# Loss of *Grin2a* Causes a Transient Delay in the Electrophysiological Maturation of Hippocampal Parvalbumin Interneurons: A Possible Mechanism for Transient Seizure Burden in Patients with Null *GRIN2A* Variants

**DOI:** 10.1101/2021.12.29.474447

**Authors:** Chad R. Camp, Anna Vlachos, Chiara Klöckner, Ilona Krey, Tue G. Banke, Nima Shariatzadeh, Sarah M Ruggiero, Peter Galer, Kristen L. Park, Adam Caccavano, Sarah Kimmel, Xiaoqing Yuan, Hongjie Yuan, Ingo Helbig, Tim A. Benke, Johannes R. Lemke, Kenneth A. Pelkey, Chris J. McBain, Stephen F. Traynelis

## Abstract

N-methyl-D-aspartate receptors (NMDARs) comprise a family of ligand-gated ionotropic glutamate receptors that mediate a calcium-permeable component to fast excitatory neurotransmission. NMDARs are heterotetrameric assemblies of two obligate GluN1 subunits (encoded by the *GRIN1* gene) and two GluN2 subunits (encoded by the *GRIN2A*-*GRIN2D* genes). Sequencing data shows that 43% (297/679) of all currently known NMDAR disease-associated genetic variants are within the *GRIN2A* gene, which encodes the GluN2A subunit. Here, we show that unlike missense *GRIN2A* variants, individuals affected with disease-associated null *GRIN2A* variants demonstrate a transient period of seizure susceptibility that begins during infancy and diminishes near adolescence. To explore this new clinical finding at that circuit and cellular level, we conducted studies using *Grin2a^+/-^* and *Grin2a^-/-^*mice at various stages during neurodevelopment. We show increased circuit excitability and CA1 pyramidal cell output in juvenile mice of both *Grin2a^+/-^* and *Grin2a^-/-^* mice. These alterations in somatic spiking are not due to global upregulation other *GRIN* genes (including *Grin2b*) nor can they be attributed to perturbations in the intrinsic excitability or action-potential firing properties of CA1 pyramidal cells. Deeper evaluation of the developing CA1 circuit led us to uncover age- and *Grin2a* gene dosing-dependent transient delays in the electrophysiological maturation programs of PV interneurons. Overall, we report that *Grin2a^+/+^* mice reach electrophysiological maturation between the neonatal and juvenile neurodevelopmental timepoints, with *Grin2a^+/-^* mice not reaching electrophysiological maturation until preadolescence, and *Grin2a^-/-^* not reaching electrophysiological maturation until adulthood. Overall, these data may represent a molecular mechanism describing the transient nature of seizure burden in disease-associated null *GRIN2A* patients.

## Introduction

N-methyl-D-aspartate receptors (NMDARs) comprise a family of ligand-gated ionotropic glutamate receptors that mediate a calcium-permeable component to fast excitatory neurotransmission (Hansen et al. 2021). NMDARs are heterotetrameric assemblies of two obligate GluN1 subunits (encoded by the *GRIN1* gene) and two GluN2 subunits (encoded by the *GRIN2A*-*GRIN2D* genes) (Hansen et al. 2021). Given their ubiquitous expression, participation in glutamatergic neurotransmission, and facilitation of calcium entry into cells, NMDARs have been implicated in a host of physiological and developmental roles including learning, memory, spatial navigation, coordinated movement, decision making, neuronal migration, morphological development, and synaptic connectivity (Collingridge 1987; Nakazawa et al. 2004; Watanabe et al. 1998; Nash and Brotchie 2000; Wang 2002; Komuro and Rakic 1993; Ultanir et al. 2007; Adesnik et al. 2008; Hansen et al. 2021).

A growing volume of sequencing data implicates genetic variation within NMDARs as a contributing factor to neuropathological conditions including epilepsy, schizophrenia, autism, intellectual disability, and developmental delay (Benke et al. 2021; Yuan et al. 2009). These genetic variants, which are absent from the healthy population, illustrate a critical role for NMDARs in basic and higher-level cognitive function (Amin, Moody, and Wollmuth 2021; Perszyk et al. 2020). Moreover, when stratified by subunit, 43% (297/679) of all currently known NMDAR disease-associated genetic variants are within the *GRIN2A* gene, which encodes the GluN2A subunit (Benke et al. 2021; Hansen et al. 2021). Neurological conditions associated with heterozygous *GRIN2A* variation present with a range of symptoms, with the most common being epilepsy and intellectual disability, coupled with some form of speech disorder, most notably oral motor apraxia (Carvill et al. 2013; Benke et al. 2021; Hansen et al. 2021). Additionally, heterozygous disease-associated variants in the *GRIN2A* gene have been identified as a high-risk factor for schizophrenia in genome-wide association studies (Singh et al. 2022; Schizophrenia Working Group of the Psychiatric Genomics 2014). Nearly one-third (98/297) of heterozygous disease-associated *GRIN2A* variants represent null variants (Hansen et al. 2021), mainly mediated by nonsense variants and deletions, in which no functional GluN2A protein would be made by the affected allele. Here, we show that unlike those with loss-of-function or gain-of-function missense *GRIN2A* variants, the majority of individuals affected with disease-associated null *GRIN2A* variants demonstrate a transient period of seizure susceptibility that begins during infancy and diminishes near adolescence.

To investigate the cellular mechanisms for this transient seizure burden, we used global *Grin2a*^+/-^ and *Grin2a*^-/-^ mice as models for null *GRIN2A* variants. While individuals with null *GRIN2A* variants are usually haploinsufficient (but see (Strehlow et al. 2022)), current data on *Grin2a*^+/-^ mice is limited. *Grin2a*^-/-^ mice, however, display neurological characteristics similar to individuals affected with null *GRIN2A* variants, such as transient cortical epileptiform discharges and deficits in spatial learning (Salmi et al. 2019; Sakimura et al. 1995). Given that this transient seizure burden manifests early in life, we hypothesized that this may be a neurodevelopmental disease, in which aberrant circuit activity during the critical plasticity period of development disrupts the excitatory-to-inhibitory balance. Loss of early GluN2A signaling would promote profound network disruptions as the GluN2B-to-GluN2A switch, a period in which the relative ratio of GluN2B:GluN2A transcript tilts in favor of GluN2A being in the majority, confers cells with faster NMDAR-mediated excitatory postsynaptic currents and less overall calcium transfer per synaptic event (Sans et al. 2000; Williams et al. 1993; Kirson and Yaari 1996; Carmignoto and Vicini 1992; Erreger et al. 2005; Schneggenburger 1996). This change in postsynaptic calcium signaling coincides with immense periods of neurodevelopment in which cells display morphological changes, synaptic connections are established and pruned, and various ion channels regulating cellular excitability are upregulated (Zhang 2004; Oswald and Reyes 2008; Piatti et al. 2011).

The GluN2A subunit is expressed in excitatory glutamatergic pyramidal cells (Perszyk et al. 2016; Hansen et al. 2021) and multiple interneuron subtypes (Perszyk et al. 2016). Inhibition or targeted knockdown of the GluN1, GluN2B, or GluN2D subunits impede interneuron development, suggesting an active role for NMDARs in interneuron function and maturation (Kelsch et al. 2014; Hanson et al. 2013; Hanson et al. 2019; Chittajallu et al. 2017). Given the transient nature of seizure susceptibility observed in null *GRIN2A* patients, the temporal expression pattern of GluN2A, and potential roles of NMDARs in circuit refinement and interneuron maturation, we hypothesized that the reduced GluN2A signaling may impact interneuron function and thereby contribute to the formation of a transiently hyperexcitable network. Our data suggest that *Grin2a^+/-^*and *Grin2a^-/-^* mice show a delay in parvalbumin-positive (PV) interneuron maturation, with resolution of aberrant interneuron function occurring at a time – post adolescence – roughly corresponding to the time null *GRIN2A* variant patients show seizure offset. These data suggest a molecular mechanism for the transient seizure burden observed in null *GRIN2A* patients and provide further evidence for GluN2A’s role in circuit maturation.

## Methods

### Animals and Breeding

All procedures involving the use of animals performed at Emory University were reviewed and approved by the Emory University IACUC, and were performed in full accordance with state and federal Animal Welfare Acts and Public Health Service policies. *Grin2a*^-/-^ mice were obtained from the laboratory of Masayoshi Mishina (University of Tokyo, Japan) and were generated via insertion of a neomycin resistance gene and Pau sequence (mRNA destabilizing and transcription-pausing signals) into the coding region of the transmembrane domain of the *Grin2a* gene as previously described (Sakimura et al. 1995). These mice were then backcrossed more than 15 times to a C57BL/6J (Jax stock number: 00664) background at Emory before use. Mice were genotyped by performing PCRs for the neomycin cassette (forward: GGGCGCCCGGTTCTT; reverse: CCTCGTCCTGCAGTTCATTCA) and the WT *Grin2a* gene (forward: GCCCGTCCAGAATCCTAAAGG; reverse: GCAAAGAAGGCCCACACTGATA). Heterozygous *Grin2a*^+/-^ mice were identified as being positive for both probes.

In order to visualize PV cells for use in electrophysiological experiments, *Grin2a*^-/-^ mice were crossed with *Pvalb*-tdTomato mice (Jax stock number: 027395 – Tg(*Pvalb*-tdTomato15Gfng)) to generate *Grin2a^+/+^*:*Pvalb*-tdTom, *Grin2a^+/-^*:*Pvalb*-tdTom, and *Grin2a^-/-^*:*Pvalb*-tdTom mice. This reporter line has already been backcrossed to a C57BL/6J background and has been previously validated to be selective and specific for PV^+^-GABAergic interneurons, including those within the CA1 subfield (Kaiser et al. 2016; Ekins et al. 2020). For identification of neonatal PV cells in acutely prepared hippocampal tissue, *Tac1*-Cre (Jax stock number: 021877 – B6;129S-*Tac1*^tm1.1(cre)Hze^/J) driver mice were crossed with eGFP-Floxed (Jax stock number: 004077 – B6;129-Gt(ROSA)26Sor^tm2Sho^/J) mice to produce *Tac1*-Cre:eGFP mice. *Tac1* has previously been described to be specifically expressed in PV cells, with no expression in MGE-derived somatostatin-positive cells (Que et al. 2021; Favuzzi et al. 2019). Additionally, since CA1 *stratum oriens* and *stratum pyramidale* had the most extensive overlap for PV and Tac1 (see Supplemental Figure S5), all recordings made from *Tac1*-positive cells were chosen in these two layers only.

The following definitions are used for mice of various ages: neonatal (P6-8), juvenile (P13-15), preadolescent (P20-26), and adult (P70-125) as described previously (Spear 2000). All mice were maintained in a conventional vivarium, given standard chow and water *ad libitum*, with a 12-hour light cycle. Both male and female mice were used in all experiments.

### Human Patient Data

Deidentified data on seizure onset and offset were obtained from consented patients under protocols approved by the University of Colorado IRB (COMIRB 16-1520), the Children’s Hospital of Philadelphia IRB, or University of Leipzig IRB (224/16-ek and 379/21-ek). Seizure offset was defined as the age of last seizure. All cases are currently ongoing.

### Gene Expression Analysis

After juvenile (P13-15) *Grin2a^+/+^*, *Grin2a^+/-^*, and *Grin2a^-/-^* mice were overdosed with inhaled isoflurane, whole hippocampi from both hemispheres were removed and placed into an Eppendorf tube and flash frozen in liquid nitrogen. A total of 18 samples were collected, with six replicates per genotype. After tissue collection, RNA was extracted using the miRNeasy Mini Kit (Qiagen; 217004) according to kit instructions. RNA quality was assessed via an Agilent 2100 Bioanalyzer and reported as RNA Integrity Numbers (RINs) in Supplemental Table . Next, multiplex mRNA expression analysis was performed using a custom designed NanoString nCounter panel consisting of all seven *GRIN* genes. Data were analyzed using NanoString’s nSolver module where samples were checked for internal quality control (QC) metrics such as imaging QC, binding density QC, positive control linearity QC, and positive control limit of detection QC (see supplemental table for details). Samples were background subtracted using negative controls (eight different hybridization probes for which no transcript has been supplied) and data that did not meet the minimum detectable threshold of 30 counts were excluded. Counts for the *Grin3b* gene were excluded from analysis for failing to meet the limit of detection. Samples were normalized to six positive hybridization controls (at the following concentrations in the 30 μL hybridization reaction: 128 fM, 32 fM, 8 fM, 2 fM, 0.5 fM, and 0.125 fM) and 11 housekeeping genes (*Aars*, *Asb10*, *Ccdc127*, *Cnot10*, *Csnk2a2*, *Fam104a*, *Gusb*, *Lars*, *Mto1*, *Supt7l*, *Tada2b*). Normalized datasets were compared for significance using the Benjamini-Yekutieli method for controlling false discovery rate.

### Acute Hippocampal Slice Preparation and Electrophysiological Recordings

After mice were overdosed with inhaled isoflurane, brains were rapidly removed and immediately placed in ice-cold ACSF (see below), and 300-µm thick horizontal, ventral hippocampal slices were made using a vibratome (Lecia, VT-1200S) in an ice-cold, sucrose-based artificial cerebrospinal fluid (aCSF) containing the following (in mM): 88 sucrose, 80 NaCl, 2.5 KCl, 1.25 HNa_2_PO_4_, 26 HNaCO_3_, 10 glucose, 2 thiourea, 3 sodium pyruvate, 5 sodium ascorbate, 12 N-acetylcysteine, 10 MgSO_4_, and 0.5 CaCl_2_ bubbled in 95% O_2_/5% CO_2_. After sectioning, slices were incubated in a sucrose-based aCSF as described above but with 4 mM MgSO_4_ at 32°C for 30 minutes then returned to room temperature for at least an hour before use. All recordings were made in the following aCSF extracellular solution (in mM): 126 NaCl, 2.5 KCl, 1.25 HNa_2_PO_4_, 26 HNaCO_3_, 20 glucose, 1.5 MgSO_4_, and 1.5 CaCl_2_ bubbled with 95% O_2_/5% CO_2_ and held at 30-32°C using an inline heater (Warner, SH-27B). Cells were visualized using an upright Olympus BX50W microscope with IR-DIC optics coupled to a Dage IR-2000 camera. Whole-cell patch clamp recordings were obtained using an Axopatch 200B (Molecular Devices) or a Multiclamp 700B (Molecular Devices, digitized at 20 kHz using a Digidata 1440a (Molecular Devices) controlled by pClamp 10.6 software (Molecular Devices). All signals were low-pass filtered at 2 kHz using a Bessel 8-pole filter (Warner, LPF-8). Patch clamp electrodes were pulled using a Sutter P1000 horizontal puller from thin-walled borosilicate capillary tubes (WPI), with a typical resistance of 4-8 MΩ.

For current clamp recordings (action-potential spiking probability, intrinsic excitability, and action potential firing properties), the following intracellular solution was used (in mM): 115 potassium gluconate, 0.6 EGTA, 2 MgCl_2_, 2 Na_2_ATP, 0.3 Na_2_GTP, 10 HEPES, 5 sodium phosphocreatine, 8 KCl, and 0.3-0.5% biocytin. After obtaining a whole-cell configuration, all cells were allowed to dialyze for 5 minutes in current-clamp mode. The liquid junction potential was not corrected and all current-clamp responses were automatically bridge-balanced using the Multiclamp 700B clamp commander software. For action-potential spiking probability experiments, CA1 pyramidal cells were held at -60 mV by injecting ± 10-30 pA of current. Any CA1 pyramidal cells that fired spontaneous action-potentials at -60 mV were rejected from analysis. A monopolar iridium-platinum stimulating electrode (FHC, Inc.) was placed in the upper 1/3^rd^ of the Schaffer collaterals to elicit a single 50 µs stimulation at a frequency of 0.03 Hz. This stimulation paradigm was used to find a stimulation intensity that would be just below a threshold that would produce a single action-potential spike of the patched CA1 pyramidal cell held at -60 mV in current-clamp mode, with a typical stimulation intensity of 40-70 µA. Once this stimulation intensity was established, the stimulation paradigm was shifted to deliver five 50 µs bursts at a frequency of 100 Hz every 30 seconds as previously described (Jami et al. 2021). A total of five epochs were recorded per cell with action-potential spiking probability calculated per stimulation number as number of spikes/5. Input resistance was calculated using the slope of voltage deflections in response to 500 ms current injections of -200, -150, -100, -50, 0, and 50 pA, with an inter-sweep interval of 2 seconds. The membrane time constant was calculated in response 20-30 sweeps of a 500 ms, -50 pA current injection, with an inter-sweep interval of 2 seconds. Composite responses were made by averaging 20-25 traces together using Clampfit (Molecular Devices). A weighted time constant was calculated using formula (1) by fitting a dual-exponential to each response in ChanneLab (Synaptosoft). Rheobase was calculated in response to 500 ms current injections starting at 0 pA and increasing by 2 pA every 2 seconds. Rheobase was defined as the minimal current injection required to elicit an action potential during the current injection period. Action potential firing frequency was calculated in response a 500 ms current injection every 2 seconds starting at -100 pA, and increasing by 20 pA for neonate and juvenile mice and 50 pA for preadolescent and adult mice, until the cell displayed depolarization-induced block of firing. Number of action potentials per current injection were calculated using pClamp (Molecular Devices). Action potential half-width, action potential amplitude, action potential threshold, and afterhyperpolarization amplitude were calculated using pClamp (Molecular Devices), from action potentials obtained during rheobase recordings.

For voltage clamp experiments (NMDAR-mediate EPSCs), the following intracellular solution was used (in mM): 100 Cs-gluconate, 5 CsCl, 0.6 EGTA, 5 BAPTA, 5 MgCl_2_, 8 NaCl, 2 Na-ATP, 0.3 Na-GTP, 40 HEPES, 5 Na-phosphocreatine, and 3 QX-314. A monopolar iridium-platinum stimulating electrode (FHC, Inc.) was placed in the upper 1/3^rd^ of the Schaffer collaterals to elicit a single 50 µs stimulation at a frequency of 0.03 Hz and the NMDAR-mediated EPSC was pharmacologically isolated with 10 μM NBQX and 10 μM gabazine. Cells were held at +40 mV and stimulation intensity was chosen to be near 50% of the maximum peak amplitude of the EPSC. A total of 8-12 epochs were recorded and averaged together. At the conclusion of recording, 200 μM DL-APV was applied to ensure responses were mediated via NMDARs. A weighted time constant was calculated using the following formula by fitting a dual-exponential function to each composite mIPSC trace in ChanneLab (Synaptosoft):

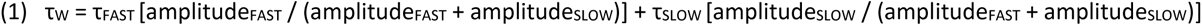

where τ_FAST_ is the fast deactivation time constant, τ_SLOW_ is the slow deactivation time constant, amplitude_FAST_ is the current amplitude of the fast deactivation component, and amplitude_SLOW_ is the current amplitude of the slow deactivation component.

For all electrophysiological recordings, series resistance was monitored throughout all experiments and was typically 8−20 MΩ. For current clamp recordings, cells were briefly switched to voltage clamp and held at -60 mV while a series of 50 ms, 5-mV square waveforms were applied to the cell. For voltage clamp recordings, this 50 ms, 5-mV square wave was included in the stimulation paradigm. Series resistance was monitored throughout the entire recording, while current clamp recordings had series resistance measurements made at the beginning and at the end of each experiment, which usually lasted 5-7 minutes total. All series resistances were measured offline by analyzing the peak of the capacitive charging spike and applying Ohm’s law. If the series resistance changed >25% during the experiment, or ever exceeded 30 MΩ, then the cell was excluded.

### Interneuron Anatomical Reconstructions

After biocytin filling during whole-cell recordings, slices were fixed with 4% PFA in 1x PBS overnight, then permeabilized with 0.3% Triton-X in 1x PBS and incubated with streptavidin Alexa546 (Invitrogen, S11225; 1:500). Resectioned slices (75-µm) were mounted on gelatin-coated slides using Mowiol mounting medium. We also found similar axonal recovery success by filling cells with 0.5% biocytin and permeabilizing slices with 1.2% Triton-X in 1x PBS for 10 minutes prior to incubation in streptavidin AlexaFluor546 without resectioning. Cells were visualized using epifluorescence microscopy and images for representative examples were obtained with confocal microscopy. Cells were reconstructed and analyzed with Sholl analysis using Neurolucida software (MBF Bioscience). Polar histograms of dendrites and axons were created using the Neurolucida function (10 degree bins). Polarity preference was determined by calculating the percentage of horizontally (150–210, 330–30 degrees) or vertically (60–120, 240–300 degrees) oriented axons for each genotype.

### Immunohistochemistry for GABAergic Interneuron Markers

After mice were overdosed with inhaled isoflurane, they were transcardially perfused with cold 1x phosphate buffered saline (PBS; pH 7.35), and subsequently perfused with cold 4% paraformaldehyde (PFA) in 1x PBS. Brains were removed and fixed for 24 hours in 4% PFA in 1X PBS before being transferred to a 30% sucrose solution dissolved in 1x PBS until the brains sank. After cryoprotection, brains were frozen in optimal cutting temperature solution (OCT, Fisher) and serial 50-µm coronal hippocampal sections were obtained, with a total of five sections per animal, spaced roughly 250-µm apart were obtained with the entire hippocampus being sampled. Slices were then transferred to a permeabilization solution containing 1.2% Triton-X in 1x PBS for 10 minutes, as previously described (Kaiser et al. 2016), before they were incubated in blocking solution containing 15% normal goat serum, 1% bovine serum albumin, and 0.5% Triton-X for 2-4 hours at room temperature. Primary antibodies were diluted in this same blocking solution at the following concentrations and incubated at 4°C for 72 hours: rabbit anti-parvalbumin (Swant, PV27; 1:5000) and rabbit anti-prepro-cholecystokinin (Frontier Institute Co., Ab-Rf350; 1:1000). After primary incubation, slices were washed 3 times in 1x PBS for 10 minutes then incubated in blocking solution containing AlexaFlour488 conjugated secondary antibodies (Abcam, ab150077; 1:1000) for 2-4 hours at room temperature. Slices were then washed 3 times in 1x PBS for 10 minutes then incubated in DAPI counterstain (Abcam, ab228549; 2 µM) for 30 minutes before being washed 3 more times in 1x PBS for 10 minutes. Slices were mounted on Superfrost Plus slides (Fisher Scientific, 12-550-15) and coverslipped with #1.5 coverslips (Thomas Scientific, 64-0717) using ProLong Gold Antifade mounting media (ThermoFisher, P36930). After mounting media had cured, slides were sealed with CoverGrip (Biotium, 23005).

### Image Acquisition and Analysis

All GABAergic interneuron marker images were acquired using a Nikon A1R HD25 line-scanning confocal microscope using NIS Elements software. The following argon laser lines were used (in nm): 405 and 488 collected using GaAsP PMTs. All images were a series of z-stacks captured using a piezo motor z-controller, with software set to acquire data in 1024 × 1024 pixel format at a 1/8^th^ frame rate dwell time. One experimenter performed all analyses and was blinded to genotype during acquisition and counting of confocal microscopy data. All GABAergic interneuron marker images were captured using Plan Apo 10x 0.45 NA objective, with some representative images captured using a Plan Apo 20x 0.75 NA objective, with images consisting of 11-14 stacks with a z-stack distance of 1 µm and a pinhole size of 19.4 µm. 4-5 hippocampal slices from each animal were imaged across 4 animals per genotype. All images were analyzed in Imaris 9.5 (Bitplane) by manually drawing hippocampal subregion boundaries and hand-counted.

### Statistical Analysis and Figure Preparation

One-way or two-way ANOVA statistical tests were performed where appropriate. For experiments where multiple statistical analyses were performed on the same dataset, our significance threshold was lowered to correct for family-wise error rate (FWER) using the Bonferroni post-hoc correction method. All studies were designed so that an effect size of at least 1 was detected at 80% or greater power. All statistical analyses were performed in Prism’s GraphPad software all figures were generated in Adobe’s Illustrator software.

## Results

### Null GRIN2A Variants May Have a Transient Seizure Burden

Despite strong selective pressure against genetic variation in *GRIN* genes, hundreds of human variants have been reported (Hansen et al. 2021). Summarizing variant data from Hansen et. al. 2021, **Figure 1A** highlights that the *GRIN2A* gene contains 44% (297/679) of all known disease-associated *GRIN* variants. **Figure 1B** shows that only 67% (199/297) of *GRIN2A* variants are missense, while the remaining 33% (98/297) are null variants as reported in (Hansen et al. 2021). Given this striking number of null *GRIN2A* variants, we utilized publicly available patient data (https://grin-portal.broadinstitute.org/) and unpublished clinical data (**Supplemental Table S1**), to evaluate seizure burden. **Figure 1C** shows disease-associated null *GRIN2A* variants display seizure onset burden at a significantly older age than disease-associated missense *GRIN2A* variants (4.5 ± 0.2 years for null *GRIN2A* variants, n=92 vs 3.1 ± 0.4 years for missense *GRIN2A* variants, n=45). We also report that 20 null *GRIN2A* patients with previous history of seizures have been seizure-free at last follow-up, with a mean seizure offset of 10.4 ± 0.8 years. These data are in stark contrast to disease-associated missense *GRIN2A* variants with available seizure onset/offset data (**Supplemental Table S2**). To date, only one disease-associated missense *GRIN2A* patient has been deemed seizure-free at the age of 1.7 years. These data suggest seizure susceptibility in disease-associated null *GRIN2A* patients may be transient, while seizure burden in disease-associated missense *GRIN2A* patients may be more permanent. More data is needed to make concrete conclusions from these findings; however, they do raise the possibility that loss of *GRIN2A* could produce a transient increase in circuit excitability.

**Figure 1.**
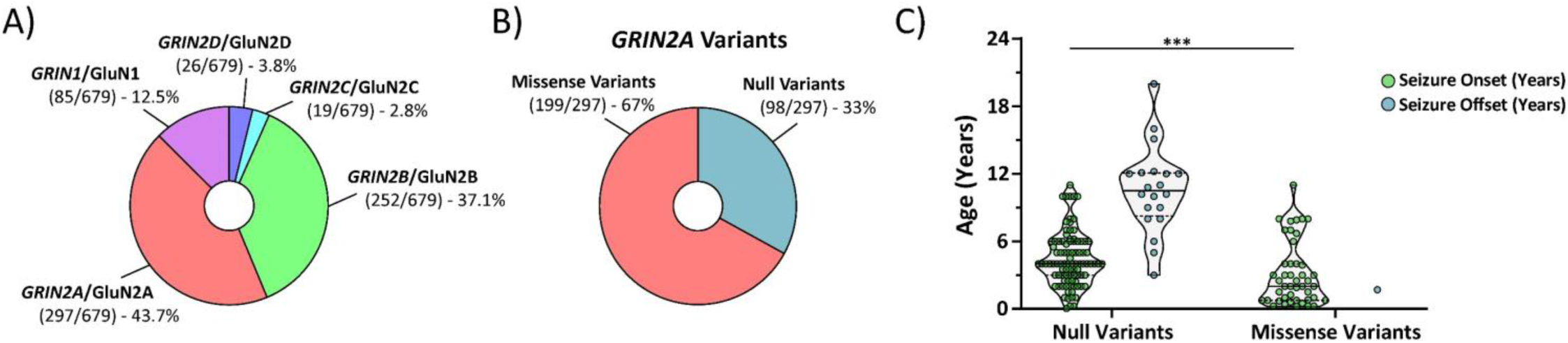
Disease-associated null *GRIN2A* patients may display a transient seizure burden not seen in disease-associated missense *GRIN2A* patients. **A)** Summary data adapted from Hansen et. al. 2021 showing that most *GRIN* variants are found in the *GRIN2A* (44 %; 297/679) gene. **B)** Summary data adapted from Hansen et. al. 2021 highlighting that 1/3^rd^ (98/297) of all *GRIN2A* variants are null variants, which include nonsense variants, as well as chromosomal insertions, deletions, inversions, and translocations. **C)** Disease-associated null *GRIN2A* variants display seizure onset burden at a significantly older age than disease-associated missense *GRIN2A* variants (4.5 ± 0.2 years for null *GRIN2A* variants, n=92 vs 3.1 ± 0.4 years for missense *GRIN2A* variants, n=45; Mann-Whitney ranked sum, p=0.0003). Currently, there are 20 disease-associated null *GRIN2A* patients with a previous history of seizures that were seizure-free at their last follow-up, with a mean seizure offset of 10.4 ± 0.8 years. These data are in stark contrast to disease-associated missense *GRIN2A* variants with available seizure offset data, as only one patient has been reported to be seizure-free at last follow-up (missense *GRIN2A* variant seizure onset age = 0.25 years with seizure offset at 1.7 years). For violin plots, solid middle line represents the median, with the dashed lines representing the 25^th^ and 75^th^ quantile. *** = p<0.001.

### Developing Hippocampus Shows Hyperexcitability in Grin2a^+/-^ and Grin2a^-/-^ Mice

Previous data using *Grin2a^-/-^* mice have shown cortical epileptiform activity in preadolescent mice (Salmi et al. 2019), as well as a prolongation of the NMDAR-mediated excitatory postsynaptic current (EPSC) onto CA1 pyramidal cells (Booker et al. 2021) and dentate gyrus granule cells (Kannangara et al. 2015). The prolongation of the NMDAR-mediated EPSC is expected given the mixed expression of rapidly deactivating GluN2A- and slowly deactivating GluN2B-containing NMDARs on excitatory pyramidal cells and the decay kinetics of a pure GluN2B-containing NMDAR population. Moreover, these data suggest that the loss of GluN2A-mediated signaling generates an increased excitability of cortical and hippocampal circuits given the longer time course of the NMDAR-mediated EPSC, however, the exact age range in when this excitability change occurs has not been explored. Since GluN2A expression shows a strong upregulation of expression during early postnatal development (Monyer et al. 1994), we explored the impact of developmental age on the prolongation of the NMDAR-mediated EPSC.

Evoked NMDAR-mediated EPSCs were obtained from CA1 pyramidal cells at various ages: neonate (P6-8), juvenile (P14-15), preadolescent (P21-26), and adult (P70+). The weighted tau describing the synaptic deactivation time course of the NMDAR-mediated EPSC was used to assess the relative ratio of GluN2A-containing NMDARs to non GluN2A-containing NMDARs as previous data have shown NMDAR complexes containing the GluN2A subunit have the fastest deactivation time of all four GluN2 subunits, both in heterologous expression systems and in native synapses (Hansen et al. 2021). The weighted tau of the NMDAR-mediated EPSC in neonatal mice was the most prolonged, regardless of genotype, suggesting very little GluN2A (**Figure 2A-2B**). The weighted tau measurement in *Grin2a^-/-^* mice was significantly more prolonged at every age when compared to *Grin2a^+/+^* mice (**Figure 2B**). These data were expected since CA1 pyramidal cells have been shown to express a mixture of GluN2A-containing and GluN2B-containing NMDARs (Hansen et al. 2014), with the presumption that GluN2B-containing NMDARs will be the only subunit present at the CA1 pyramidal cell synapse in the *Grin2a^-/-^*mouse. The weighted tau of the NMDAR-mediated EPSCs in juvenile and preadolescent mice were significantly different across all three genotypes, with each following a *Grin2a* gene dosing-dependent shortening of weighted tau (**Figure 2B**). In adult mice, the weighted tau measurements in *Grin2a^+/+^* and *Grin2a^+/-^* mice overlap, suggesting that *Grin2a^+/-^* mice eventually reach a wildtype weighted tau. The increased weighted tau between the preadolescent and adult timepoints in *Grin2a^+/+^* mice was unexpected and could be due to an increased insertion of GluN2B-containing NMDARs in adult mice after the preadolescent critical period of plasticity and development. More importantly, these data highlight that the juvenile developmental time point may be the start of aberrant synaptic signaling, at least in terms of charge transfer and calcium influx.

**Figure 2.**
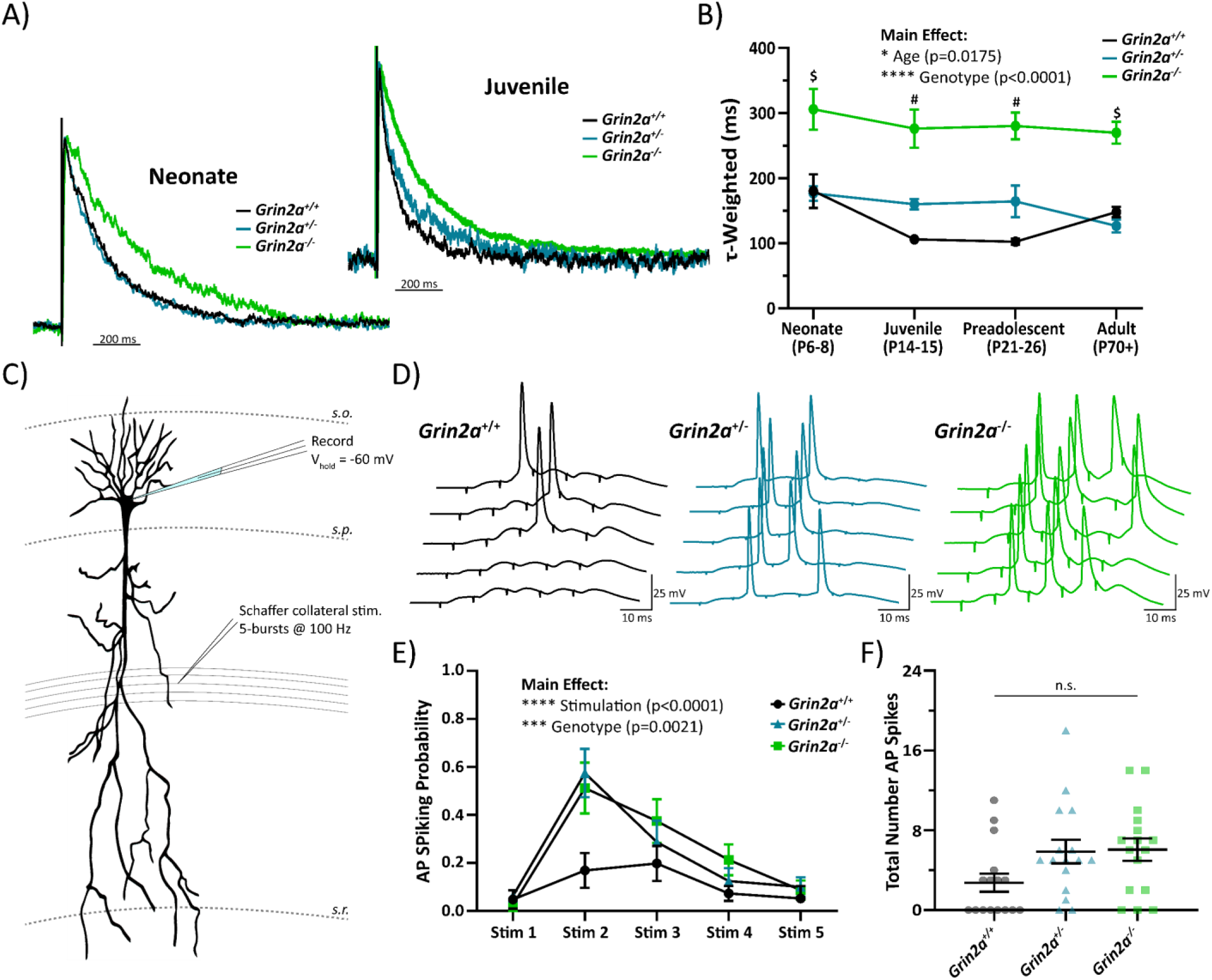
Juvenile CA1 circuit shows hyperexcitability in *Grin2a^+/-^* and *Grin2a^-/-^*mice. **A)** Evoked NMDAR-mediated excitatory postsynaptic currents (EPSCs) onto CA1 pyramidal cells from *Grin2a^+/+^, Grin2a^+/-^*, *Grin2a^-/-^* at various ages during development, with normalized representative traces shown for neonate and juvenile mice. **B)** Two-way ANOVA showed significant main effects for both age (F_3, 93_ = 3.55; p = 0.0175) and genotype (F_3, 93_ = 71.4; p<0.0001) of the tau-weighted describing the NMDAR-mediated EPSC decay time. At both neonate and adult timepoints, only *Grin2a^+/+^* and *Grin2a^-/-^* mice have significantly different NMDAR-mediated EPSC decay times, whereas at the juvenile and preadolescent timepoints, all three genotypes have significantly different NMDAR-mediated EPSC decay times. Given that the juvenile stage was the earliest developmental window with separation at all three genotypes for the NMDAR-mediated EPSC decay time, we conducted further experiments at this age. **C)** CA1 pyramidal cells from juvenile mice were current clamped at -60 mV and Schaffer collateral afferents were stimulated five times at 100 Hz for a total of 5 epochs. Stimulation intensity was set just below threshold to produce an action-potential spike after a single Schaffer collateral stimulation. **D)** Representative traces showing action-potential spiking in response to successive Schaffer collateral stimulations. **E)** Action-potential spiking probability for each stimulus averaged over 5 epochs across all genotypes. Two-way ANOVA showed significant main effects for both genotype (F_2, 225_ = 6.351; p = 0.0021) and stimulation number (F_4, 225_ = 16.82; p < 0.0001), however, there was no interaction. **F)** Total number of action potentials elicited over 5 epochs of 5-burst Schaffer collateral stimulation. One-way ANOVA showed there was no significant difference across the three genotypes (F = 2.997; p = 0.06). Data represented show mean ± SEM. AP = action-potential; *s.o.* = *stratum oriens*; *s.p.* = *stratum pyramidale*; *s.r.* = *stratum radiatum*; n.s. = not significant. $ = *Grin2a^-/-^* significantly different than both *Grin2a^+/+^* and *Grin2a^+/-^* at that age; # = all three genotypes significantly different than each other at that age.

Since the earliest divergence in NMDAR-mediated EPSC weighted tau across all three genotypes occurred in juvenile mice, this age was our benchmark to explore what other changes to hippocampal circuit function, if any, occur in *Grin2a^+/-^* and *Grin2a^-/-^* mice. Thus, we recorded action-potential generation probability in juvenile CA1 pyramidal cells during five Schaffer collateral stimulations at 100 Hz (e.g. a 50 ms burst of 5 pulses) to assess circuit output after excitatory afferent signaling. Two-way ANOVA shows a statistically significant main effect for both stimulation number and genotype (**Figure 2E; Supplemental Table S3**), however, there are no significant interactions nor differences in the total number of action-potentials generated (**Figure 2E-2F**; **Supplemental Table S3**). These data suggest that *Grin2a^+/-^* and *Grin2a^-/-^* mice are more likely to fire action-potentials when subjected to similar excitatory afferent activity compared to *Grin2a^+/+^* mice.

One possible interpretation of these data is that the loss of *Grin2a* triggers a compensatory upregulation of other NMDAR subunits, especially GluN2B. Data from whole hippocampus mRNA screening indicates no significant differences in the *Grin1*, *Grin2b*, and *Grin2c* genes across all three genotypes of juvenile mice (**Supplemental Figure S1**). mRNA data do show a statistically significant upregulation of both *Grin2d* and *Grin3a*, but these upregulations only occur in *Grin2a^-/-^* mice (**Supplemental Figure S1**). Since our synaptic spiking data show similar hyperexcitability in both *Grin2a^+/-^* and *Grin2a^-/-^* mice, these data are likely not explanatory for our observed spiking phenotype. Alternatively, heightened synaptic spiking could be due to alterations in the intrinsic excitability of the CA1 pyramidal cells themselves. We show, however, no differences in any measurable electrophysiological passive property such as resting membrane potential or input resistance, as well as no changes in action-potential firing properties (**Supplemental Figure S2**; **Supplemental Table S4**). Thus, changes in action-potential firing probability in *Grin2a^+/-^* and *Grin2a^-/-^* mice are not due to gross alterations in NMDAR subunit mRNA expression or in CA1 pyramidal cell intrinsic excitability. In addition to providing direct excitatory afferent signaling onto pyramidal cells, Schaffer collateral stimulation will also provide excitatory tone onto feedforward GABAergic interneurons. Additionally, we show that the total loss of the *Grin2a* gene promotes the upregulation of *Grin2d* mRNA, which has been found to be exclusive to GABAergic interneurons in the hippocampus (Perszyk et al. 2016; von Engelhardt et al. 2015). For these reasons, we chose to further explore the GABAergic interneuron network in developing CA1.

### Alterations in Hippocampal PV Cell Density

Alterations in synaptic excitability as described Figure 2 could be due to changes in several features of the hippocampal circuit, including GABAergic inhibition which is mediated by a wide range of different interneurons. Inhibitory GABAergic basket cells primarily make somatic inhibitory synaptic connections where they exert profound control on pyramidal cell firing (Veres, Nagy, and Hajos 2017), and in CA1, are heavily innervated by Schaffer collateral afferents (Pelkey et al. 2017). CA1 basket cells can be split into two main subtypes: parvalbumin (PV)-expressing cells and cholecystokinin (CCK)-expressing cells (Pelkey et al. 2017). PV and CCK cells have opposing transcriptomic profiles, and each represents a major subtype of interneuron arising from the medial ganglionic eminence (MGE) and caudal ganglionic eminence (CGE), respectively (Pelkey et al. 2017). Since early pyramidal cell activity has been shown to control interneuron apoptosis (Wong et al. 2018), and we report alterations to the NMDAR-mediated EPSC onto young pyramidal cells, we hypothesized that the loss of *Grin2a* may impact basket cell density in CA1. We therefore determined cell density of PV and CCK interneurons in *Grin2a*^+/+^, *Grin2a*^+/-^, and *Grin2a*^-/-^ mice via immunohistochemical staining.

The total loss of *Grin2a* promotes an upregulation of PV cells in CA1 (**Figure 3A-3B; Supplemental Table S5**) compared to both *Grin2a*^+/+^ and *Grin2a*^+/-^mice. Although the overall cell density of PV cells is altered in *Grin2a*^-/-^ mice, there is no difference in PV cellular lamination (**Figure 3A** and **3C; Table Supplemental S5**) regardless of genotype, with most PV cells residing in *stratum oriens* and *stratum pyramidale* as previously reported (Pelkey et al. 2017). Unlike PV cells, we show that the loss of *Grin2a* does not impact CCK cell density (**Figure 4A-4B; Table Supplemental S6**) or CCK cellular lamination (**Figure 4A** and **4C; Supplemental Table S6**). Given that we observed an increase in CA1 PV cell density in *Grin2a^-/-^*mice, we also wanted to check if these extra PV cells showed any morphological differences from age-matched CA1 PV cells from *Grin2a^+/+^*mice. Using biocytin-backfilled reconstructions of preadolescent *Grin2a^+/+^*and *Grin2a^-/-^* CA1 PV cells, we report no change in dendritic/axonal morphology and laminar targeting (**Supplemental Figure S3**). Thus, these data indicate that the loss of *Grin2a* may preferentially impact PV interneuron survival/apoptosis in CA1. Moreover, the effect of increased cell density in only *Grin2a*^-/-^ mice suggests that a threshold of aberrant pyramidal cell activity must be reached before PV cell survival/apoptosis is impacted, with little to no impact on cellular morphology or neurite targeting. Pyramidal cell activity thus far has only been shown to impact MGE-derived interneurons, with no data on survival/apoptosis of CGE-derived interneurons (Wong et al. 2018). Moreover, CCK cells may not be affected since previous reports have shown that these cells have little to no GluN2A-mediated synaptic signaling (Matta et al. 2013; Booker et al. 2021). Although a change in the PV cell density will likely contribute to aberrant CA1 circuit function, these data alone are unlikely to explain our observed action potential spiking phenotype.

**Figure 3.**
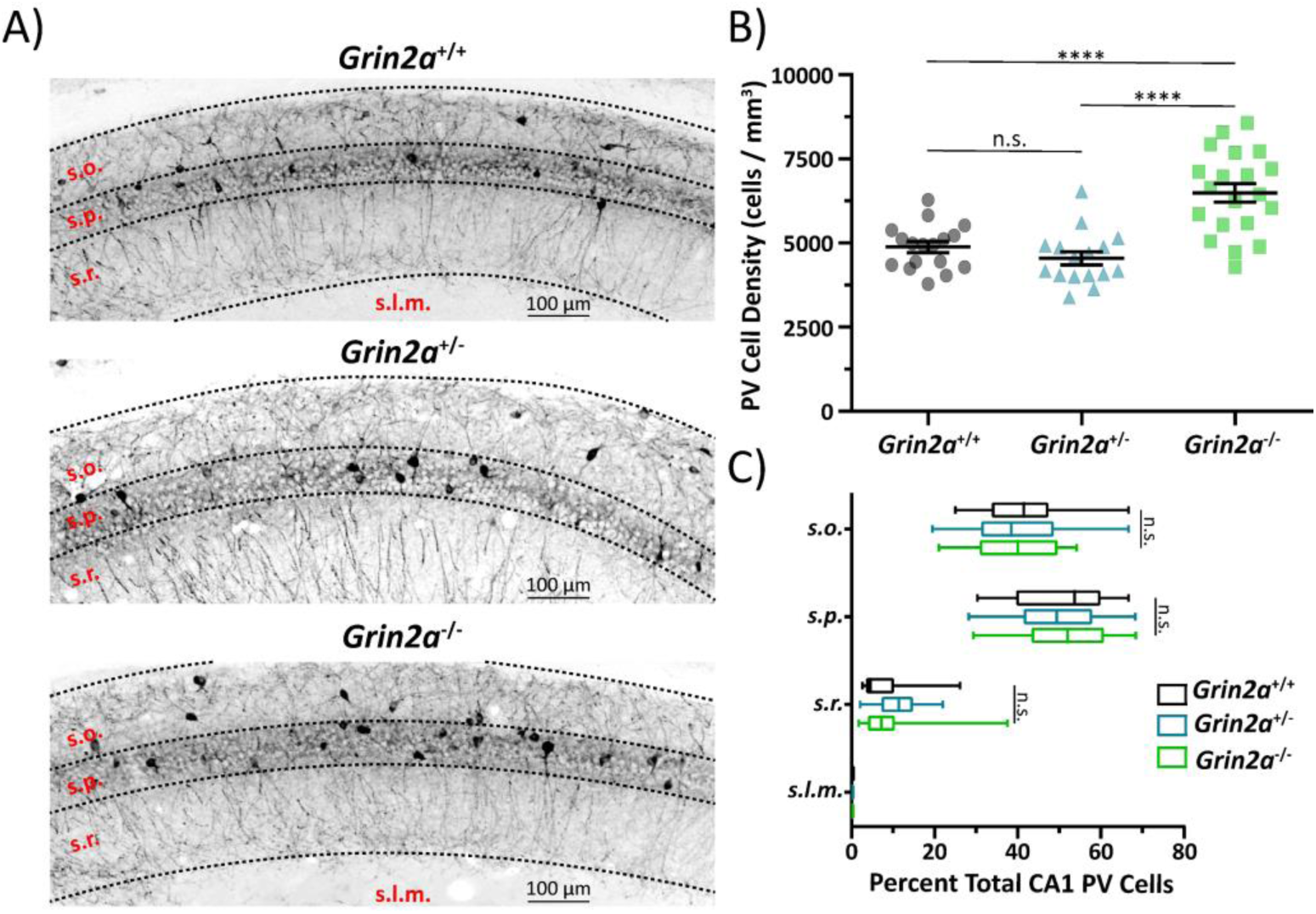
Loss of *Grin2a* causes an increase in parvalbumin (PV) cell density in CA1. **A)** Representative images of CA1 hippocampal sections stained for PV in preadolescent mice. **B)** CA1 PV cell density is significantly increased in *Grin2a*^-/-^ mice compared to *Grin2a*^+/+^ mice (6,488 ± 276 cells per mm^3^ in *Grin2a*^-/-^ vs 4,875 ± 162 cells per mm^3^ in *Grin2a*^+/+^; one-way ANOVA, p<0.0001) and *Grin2a*^+/-^ mice (6,488 ± 276 cells per mm^3^ in *Grin2a*^-/-^ vs 4,542 ± 198 cells per mm^3^ in *Grin2a*^+/-^; one-way ANOVA, p<0.0001). There is no difference in CA1 PV cell density between *Grin2a*^+/+^ and *Grin2a*^+/-^ mice (one-way ANOVA, p=0.58). **C)** Despite an increase in cell density in *Grin2a*^-/-^ mice, there is no difference in PV CA1 cellular lamination across all three genotypes. Data represented show mean ± SEM. *s.o.* = *stratum oriens*; *s.p.* = *stratum pyramidale*; *s.r.* = *stratum radiatum*; *s.l.m.* = *stratum lacunosum moleculare*; PV = parvalbumin; **** = p<0.0001; n.s. = not significant.

**Figure 4.**
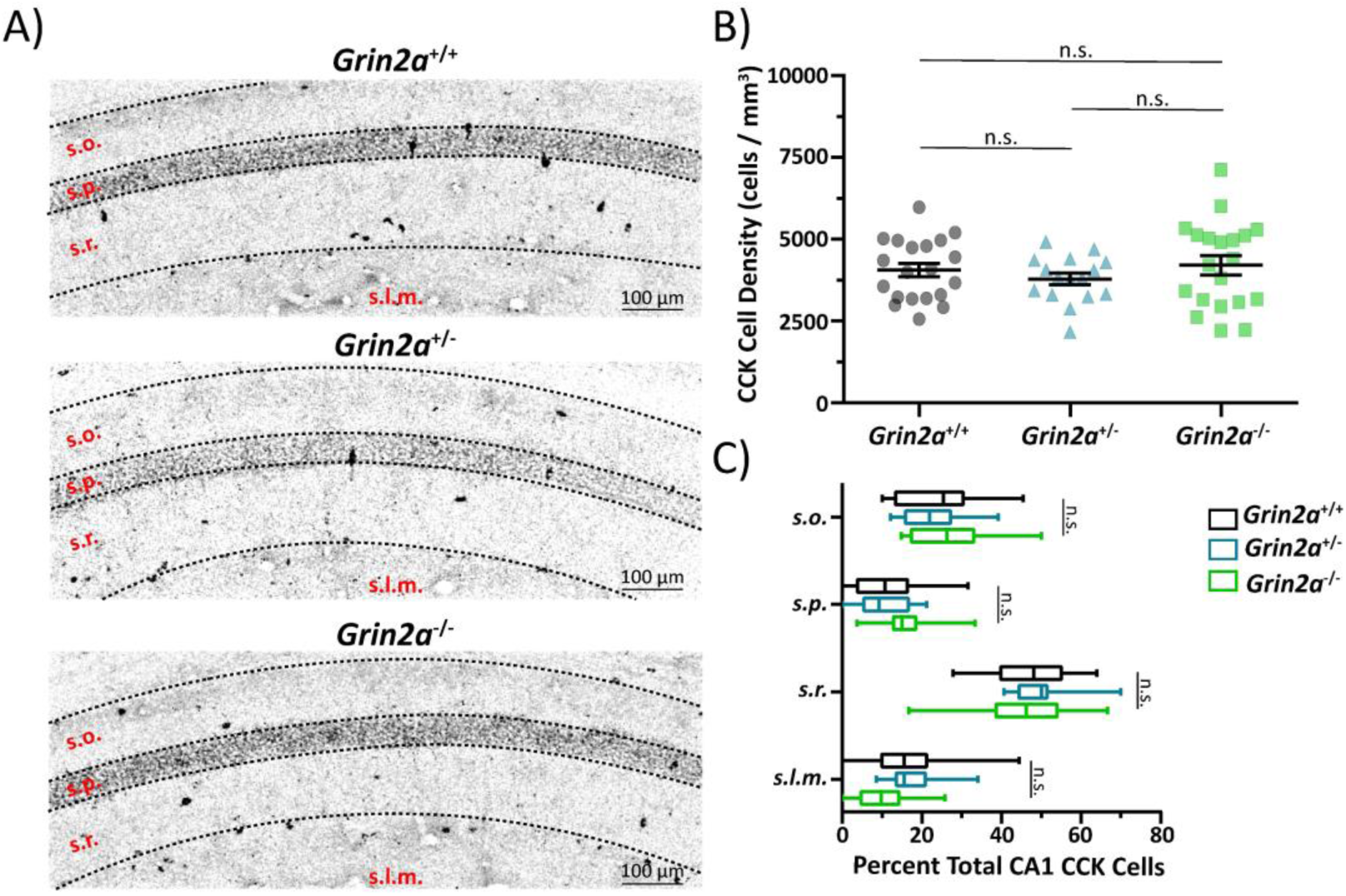
Loss of *Grin2a* does not alter cholecystokinin (CCK) cell density in CA1. **A)** Representative images of CA1 hippocampal sections stained for CCK in preadolescent mice. **B)** CA1 CCK cell density is unchanged across all three genotypes (one-way ANOVA, p=0.49). **C)** There is no difference in CCK CA1 cellular lamination across all three genotypes. Data represented show mean ± SEM. *s.o.* = *stratum oriens*; *s.p.* = *stratum pyramidale*; *s.r.* = *stratum radiatum*; *s.l.m.* = *stratum lacunosum moleculare*; CCK = cholecystokinin; n.s. = not significant.

### Age-Dependent Changes in Passive and Action-Potential Firing Properties of CA1 PV Cells

In addition to controlling interneuron survival/apoptosis, local pyramidal cell activity has also been implicated in controlling the maturation of GABAergic interneurons (Wong et al. 2018; Lim et al. 2018). Previous work has shown that PV cells undergo an electrophysiological maturation of both their passive and action-potential firing properties, however, many of these studies have been performed in cortical PV cells (Goldberg et al. 2011; Miyamae et al. 2017; Okaty et al. 2009) and dentate gyrus PV cells (Doischer et al. 2008). Since clear data highlighting an electrophysiological maturation in CA1 is limited (however, see (Que et al. 2021)), we first wanted to demonstrate that hippocampal PV cells also show an electrophysiological maturation pattern like those seen in cortex.

Detailed analysis of the electrophysiological maturation pattern in young PV cells has been hampered by the age-dependent expression of the PV gene itself, which begins around P14 (de Lecea, del Rio, and Soriano 1995). To bypass this limitation, we used another driver mouse line, *Tac1*-Cre (Jax #:021877), to obtain neonatal (P6-8) PV cells recordings in developing hippocampus. *Tac1* is highly expressed in PV cells, with little to no expression reported in MGE-derived somatostatin cells (Que et al. 2021; Favuzzi et al. 2019). Additionally, *Tac1* expression begins early in embryogenesis and is sustained well into adulthood (Allen Mouse Brain Atlas). To confirm that *Tac1*-Cre mice are a viable tool for studying neonatal PV cells, we first performed immunohistochemical colocalization staining to confirm that PV cells do indeed express the *Tac1* gene. Adult tissue from *Tac1*-Cre × Floxed-eGFP mice were stained for eGFP and PV, with data reporting that 90 ± 5% of all PV-positive cells also stained positive for eGFP (**Supplemental Figure S4**), whereas 52 ± 3% of eGFP positive cells also stained positive for PV (**Supplemental Figure S4**). Closer evaluation of these data reveal that the majority of PV/eGFP overlap occurs in the *stratum oriens* and *stratum pyramidale* layers of CA1. For these reasons, only *Tac1*-positive cells from these two laminae were chosen for recording. Moreover, all patch-clamped cells using *Tac1*-Cre × Floxed-eGFP mice were biocytin-backfilled and visually confirmed to have no dendritic spines, as some *Tac1*-positive puncta appear to be pyramidal cells (**Supplemental Figure S5**). We also compared several passive and action-potential firing properties in juvenile (P14-15) *Tac1*-Cre positive cells against a traditional *Pvalb*-TdTomato driver line (Jax #:027395) and found no differences (**Supplemental Figure S5**).

Using *Tac1*-Cre mice for neonatal recordings and *Pvalb*-TdTomato mice for juvenile, preadolescent, and adult recordings we assayed various passive and action-potential firing properties of CA1 PV cells at four stages during neurodevelopment. We report that neonatal CA1 PV cells show a transient prolongation of their membrane time constant, which is significantly increased compared to juvenile, preadolescent, and adult CA1 PV cells (**Figure 5B-5C; Supplemental Table S7**). A similar trend is observed for input resistance in neonatal CA1 PV Cells (**Supplemental Table S7**). The action-potential half-width of neonatal CA1 PV cells is also transiently prolonged and is significantly increased compared to juvenile, preadolescent, and adult cells (**Figure 5D-5E; Supplemental Table S7**). A transient prolongation of the membrane time constant and action-potential half-width observed here matches previously reported data for developing cortical PV cells (Goldberg et al. 2011). We also report dampening of the maximum action-potential firing frequency in developing CA1 PV cells. Neonatal CA1 PV cells maximum action-potential firing capacity is significantly decreased compared to juvenile, preadolescent, and adult CA1 PV cells (**Figure 5F-5G; Supplemental Table S7**). The maximum action-potential firing frequency in juvenile CA1 PV cells is also significantly decreased compared to adult cells (**Figure 5F-5G; Supplemental Table S7**). An age-dependent increase in the fast-spiking nature of PV cells has been well characterized and is thought to be driven by a delay in the upregulation of the rapid Kv3-family of voltage-gated potassium channels (Goldberg et al. 2011). The current required for depolarization-induced block of action-potential firing in neonatal CA1 PV cells is significantly decreased compared to juvenile, preadolescent, and adult cells (**Figure 5F and 5H; Supplemental Table S7**). Here, the decreased current for depolarization-induced block is likely caused by an increased input resistance measured in neonatal mice. In all, we show that CA1 PV cells do undergo significant electrophysiological maturation programs, transforming them from simple signal propagators to precise signaling processors.

**Figure 5.**
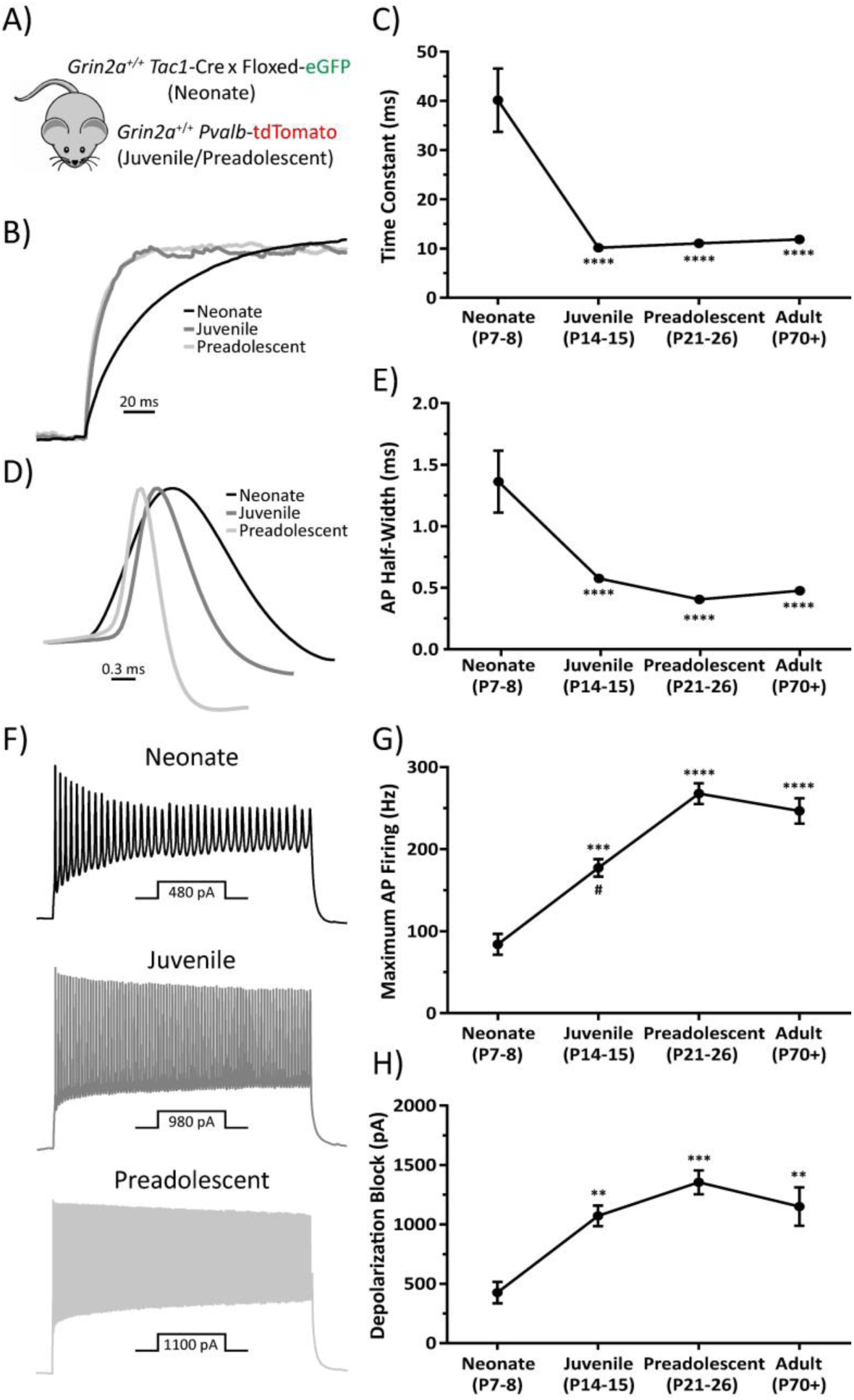
CA1 PV cells undergo electrophysiological maturation of passive and action-potential firing properties. **A)** PV cells were visualized using either *Pvalb*-TdTomato or *Tac1*-Cre × floxed eGFP (See Supplemental Figures S5 and S6). **B)** Representative, amplitude-normalized repolarization traces to highlight differences in membrane time constant following a -50 pA current injection at different developmental timepoints. **C)** Membrane time constant is significantly prolonged in neonatal mice (F = 40.68, one-way ANOVA; p<0.0001 for post-hoc multiple comparisons with every other developmental timepoint). **D)** Action-potential half-width is significantly prolonged in neonatal mice (F = 28, one-way ANOVA; p<0.0001 for post-hoc multiple comparisons with every other developmental timepoint). **E)** Representative, amplitude-normalized single action-potential traces to highlight differences in half-width at different developmental timepoints. **F)** Representative action-potential trains elicited by various current injections depicted below each train to illustrate change in maximum action-potential firing frequency during development. Traces shown are those just below threshold for depolarization-induced block of action-potential firing. **G)** Maximum action-potential firing frequency is significantly decreased in neonatal mice (F = 28.3, one-way ANOVA; p<0.001 for post-hoc multiple comparison test with juvenile mice, and p<0.0001 for post-hoc multiple comparison test with preadolescent and adult mice). Juvenile mice also show a significantly decreased maximum action-potential firing frequency compared to preadolescent mice (one-way ANOVA post-hoc multiple comparison test; p=0.0039). **F)** Current required for depolarization-induced block of action-potential firing is significantly decreased in neonatal mice (F = 8.36, one-way ANOVA; p<0.01 for post-hoc multiple comparison test with juvenile and adult mice, and p<0.0001 for post-hoc multiple comparison test with preadolescent mice). Symbols are mean ± SEM. AP = action-potential; depolarization block = Current required for depolarization-induced block of action-potential firing. ** = p<0.01; ***p<0.001; **** = p<0.0001; # = juvenile mice significantly different than adult mice in one-way ANOVA post-hoc multiple comparison test.

### Electrophysiological Maturation of PV Cells is Delayed in Grin2a^+/-^ and Grin2a^-/-^ Mice

After showing that CA1 PV cells in *Grin2a^+/+^* mice undergo electrophysiological maturation of their passive and action-potential firing properties, we tested our hypothesis that altered network activity may impact the rate of PV cell maturation in *Grin2a^+/-^*and *Grin2a^-/-^* mice. We generated *Grin2a^+/+^*:*Pvalb*-tdTom, *Grin2a^+/-^*:*Pvalb*-tdTom, and *Grin2a^-/-^*:*Pvalb*-tdTom mice via selective breeding (see methods) to visualize PV cells in CA1 across these three genotypes. We found no difference in the resting membrane potential of juvenile, preadolescent, or adult CA1 PV cells in *Grin2a^+/+^*, *Grin2a^+/-^*, and *Grin2a^-/-^* mice (**Figure 6B; Supplemental Table S8**). We did not find any statistically significant main effects of age or genotype for PV cellular capacitance (**Figure 6C**; **Supplemental Table S8**). Both the membrane time constant and input resistance both had statistically significant main effects for age and genotype, indicating a transient increase in both measures of passive membrane excitability (**Figure 6E-6F**; **Supplemental Table S8**). Moreover, we found statistically significant interactions for membrane time constant values indicating that membrane time constant decreases in an age- and *Grin2a* gene-dosing dependent manner (**Figure 6D-6E**; **Supplemental Table S8**). Overall, we found that *Grin2a^+/-^* mice didn’t reach *Grin2a^+/+^*membrane time constant values until preadolescence, while *Grin2a^-/-^*mice didn’t reach *Grin2a^+/+^* membrane time constant values until adulthood. Importantly, though, all membrane time constant values eventually did attain *Grin2a^+/+^* levels (**Figure 6D-6E**; **Supplemental Table S8**). We saw this same trend for input resistance values (**Figure 6F**; **Supplemental Table S8**). The sum of these data illustrates that there is an age- and genotype-dependent transient delay in the passive membrane excitability of CA1 PV cells. That is, values reach those of *Grin2a^+/+^*eventually, but remain at immature levels for an extended period that is dependent on the number of functional copies of *Grin2a*.

**Figure 6.**
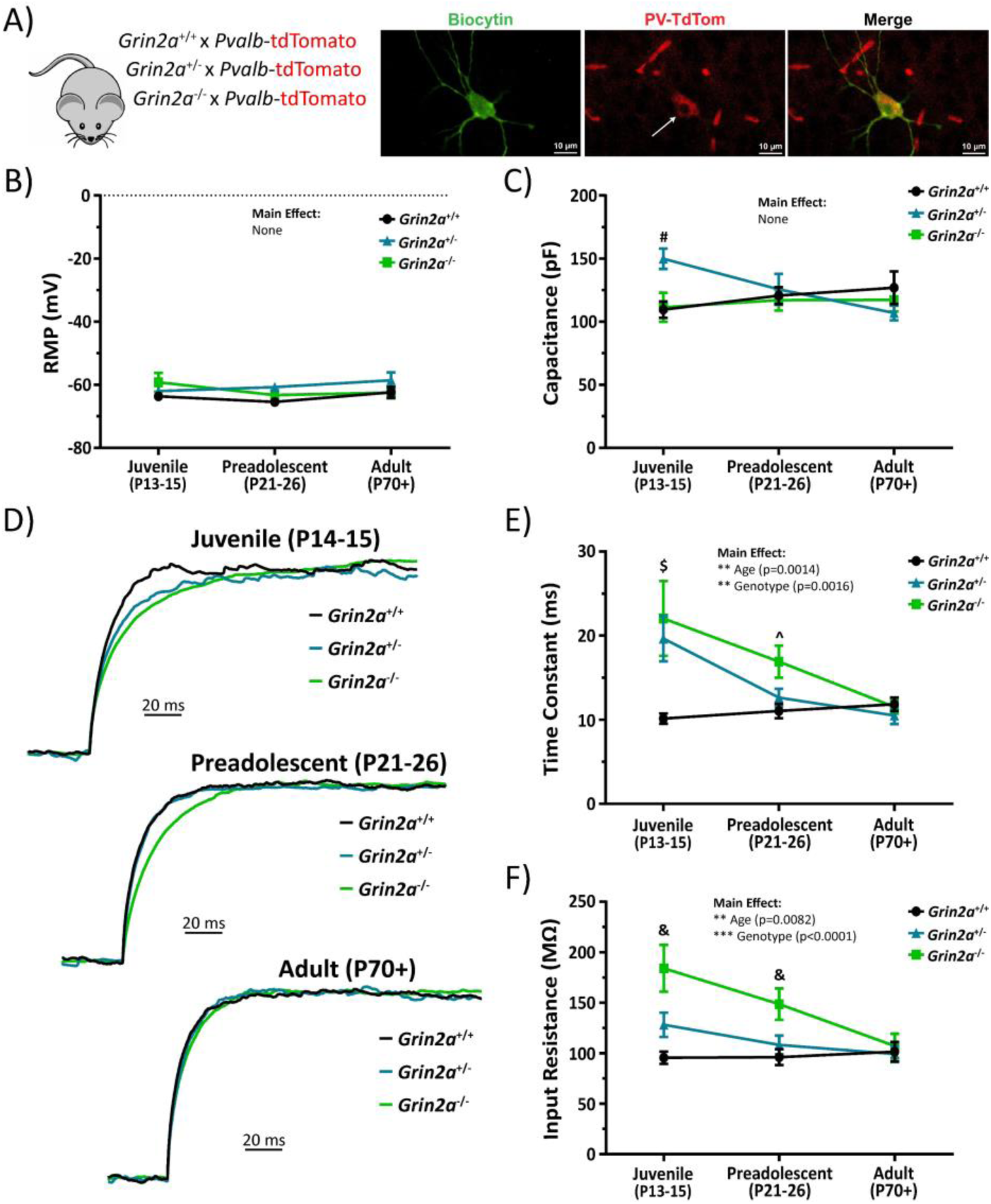
The loss of *Grin2a* causes a transient change in passive electrophysiological properties in CA1 PV cells. **A)** Mouse model used to visualize PV cells in CA1 next to examples of a biocytin backfilled CA1 PV cell that was stained for TdTomato to indicate successful cellular identification. **B)** No change in resting membrane potential across all three genotypes at different developmental time points (two-way ANOVA). **C)** No main effect of cellular capacitance across genotype (F_2, 136_ = 1.3; p=0.28; two-way ANOVA) or age (F_2, 136_ = 0.32; p=0.72; two-way ANOVA), however, there is a statistically significant interaction such that juvenile *Grin2a^+/-^*mice have a higher cellular capacitance than both *Grin2a^+/+^* mice (150 ± 8.0 pF for *Grin2a^+/-^* vs 110 ± 6.5 pF for *Grin2a^+/+^*; p=0.0061) and *Grin2a^-/-^* mice (150 ± 8.0 pF for *Grin2a^+/-^* vs 111 ± 11 pF for *Grin2a^-/-^*; p=0.014). **D)** Representative, amplitude-normalized repolarization traces to highlight differences in membrane time constant following a -50 pA current injection at different developmental timepoints. **E)** Membrane time constant measurements show statistically significant main effects for both age (F_2, 136_ = 6.9; p=0.0014; two-way ANOVA) and genotype (F_2, 136_ = 6.8; p=0.0016; two-way ANOVA). There are also several statistically significant interactions such that both juvenile *Grin2a^+/-^* mice (20 ± 2.7 ms for *Grin2a^+/-^* vs 10 ± 0.6 ms for *Grin2a^+/+^*; p=0.0011) and *Grin2a^-/-^* mice (22 ± 4.4 ms for *Grin2a^-/-^*vs 10 ± 0.6 ms for *Grin2a^+/+^*; p<0.0001) displayed higher membrane time constants than *Grin2a^+/+^* mice. Additionally, preadolescent *Grin2a^-/-^* mice showed a higher membrane time constant than *Grin2a^+/+^* mice (17 ± 1.9 ms for *Grin2a^-/-^* vs 11 ± 0.9 ms for *Grin2a^+/+^*; p=0.048). **F)** Input resistance measurements show statistically significant main effects for both age (F_2, 133_ = 5.0; p=0.0082; two-way ANOVA) and genotype (F_2, 133_ = 12.2; p<0.0001; two-way ANOVA). There are also several statistically significant interactions such that juvenile *Grin2a^-/-^* mice displayed higher input resistances than *Grin2a^+/+^* mice (184 ± 23 MΩ for *Grin2a^-/-^* vs 96 ± 6.1 MΩ for *Grin2a^+/+^*; p<0.0001) and *Grin2a^+/-^*mice (184 ± 23 MΩ for *Grin2a^-/-^* vs 128 ± 12 MΩ for *Grin2a^+/+^*; p=0.0054). Additionally, preadolescent *Grin2a^-/-^* mice showed a higher input resistance than *Grin2a^+/+^* mice (149 ± 15 MΩ for *Grin2a^-/-^* vs 96 ± 7.9 MΩ for *Grin2a^+/+^*; p=0.0031) and *Grin2a^+/-^* mice (149 ± 15 MΩ for *Grin2a^-/-^* vs 108 ± 9.3 MΩ for *Grin2a^+/+^*; p=0.043). The sum of these data indicates an age- and gene-dependent transient delay in various passive electrical properties of CA1 PV cells. Symbols are mean ± SEM. RMP = resting membrane potential; # = *Grin2a^+/-^* significantly different than both *Grin2a^+/+^* and *Grin2a^-/-^* at that age; $ = *Grin2a^+/+^* significantly different than both *Grin2a^+/-^* and *Grin2a^-/-^* at that age; ^ = *Grin2a^-/-^* significantly different than *Grin2a^+/+^* at that age; & = *Grin2a^-/-^* significantly different than both *Grin2a^+/+^* and *Grin2a^+/-^* at that age; ** = p<0.01; ***p<0.001.

We also examined the action-potential waveform and firing properties of CA1 PV cells in *Grin2a^+/-^* and *Grin2a^-/-^*mice during development. We show no statistically significant effects of age or genotype on rheobase or action-potential amplitude, as well as no significant interactions for each measure (**Figure 7A-7B**; **Supplemental Table S9**). Action-potential half-width, however, shows statistically significant effects for both age and genotype (**Figure 7C-7D**; **Supplemental Table S9**). Statistically significant interactions for the action-potential half-width indicate that action-potential half-width decreases in an age- and *Grin2a* gene-dosing dependent manner (**Figure 7C-7D**; **Supplemental Table S9**). Overall, we found that *Grin2a^+/-^*mice didn’t reach *Grin2a^+/+^* action-potential half-width values until preadolescence, while *Grin2a^-/-^* mice didn’t reach *Grin2a^+/+^* action-potential half-width values until adulthood. Importantly, though, all action-potential half-width values eventually did attain *Grin2a^+/+^* levels (**Figure 7D-7E**; **Supplemental Table S9**). These data indicate a significant but transient prolongation of the action-potential half-width that is dependent on both age and genotype, in line with data obtained on the membrane time constant and input resistance. The afterhyperpolarization amplitude shows a statistically significant main effect for age, but not for genotype, which suggests that the channels responsible for afterhyperpolarization amplitude are not subjected to the same delay observed for other measures like membrane time constant, input resistance, and action-potential half-width (**Figure 7E**; **Supplemental Table S9**). We also found that both the maximum action-potential firing frequency before depolarization-induced block and the current required to reach depolarization-induced block of action potential firing show statistically significant main effects for age and genotype (**Figure 8A-8E**; **Supplemental Table S9**).

**Figure 7.**
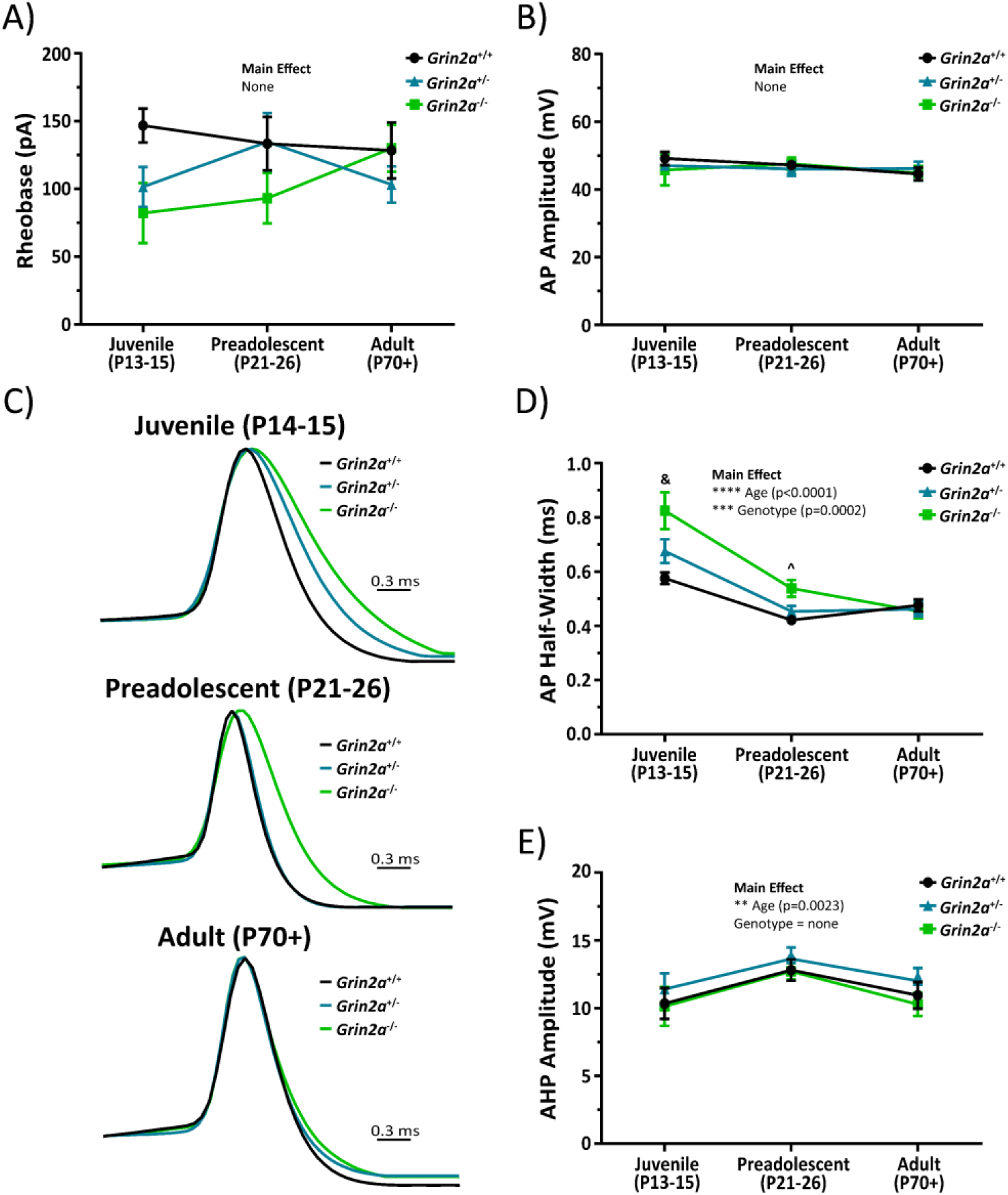
The loss of *Grin2a* causes a transient change in action-potential waveform properties of CA1 PV cells. There are no differences in **A)** rheobase or **B)** action-potential amplitude regardless of age or genotype in CA1 PV cells. **C)** Representative, amplitude-normalized single action-potential traces to highlight differences in half-width at different developmental timepoints. **D)** Action-potential half-width measurements show statistically significant main effects for both age (F_2, 134_ = 47; p<0.0001; two-way ANOVA) and genotype (F_2, 134_ = 9.3; p=0.0002; two-way ANOVA). There are also several statistically significant interactions such that both juvenile *Grin2a^+/+^* mice (0.58 ± 0.02 ms for *Grin2a^+/+^* vs 0.82 ± 0.07 ms for *Grin2a^-/-^*; p<0.0001) and *Grin2a^+/-^* mice (0.68 ± 0.04 ms for *Grin2a^+/-^*vs 0.82 ± 0.07 for *Grin2a^-/-^*; p<0.005) displayed longer action-potential half-widths than *Grin2a^-/-^* mice. Additionally, preadolescent *Grin2a^-/-^* mice showed longer action-potential half-widths than *Grin2a^+/+^* mice (0.54 ± 0.03 ms for *Grin2a^-/-^* vs 0.42 ± 0.01 ms for *Grin2a^+/+^*; p=0.0152). **E)** Afterhyperpolarization amplitude of the action-potential waveform showed a significant main effect for age (F_2, 127_ = 6.36; p=0.0023; two-way ANOVA), but no main effect for genotype (F_2, 127_ = 2; p=0.14; two-way ANOVA). Symbols are mean ± SEM. AP = action-potential; AHP = afterhyperpolarization; ^ = *Grin2a^-/-^*significantly different than *Grin2a^+/+^* at that age; & = *Grin2a^-/-^* significantly different than both *Grin2a^+/+^*and *Grin2a^+/-^* at that age; ** = p<0.01; ***p<0.001; **** = p<0.0001.

**Figure 8.**
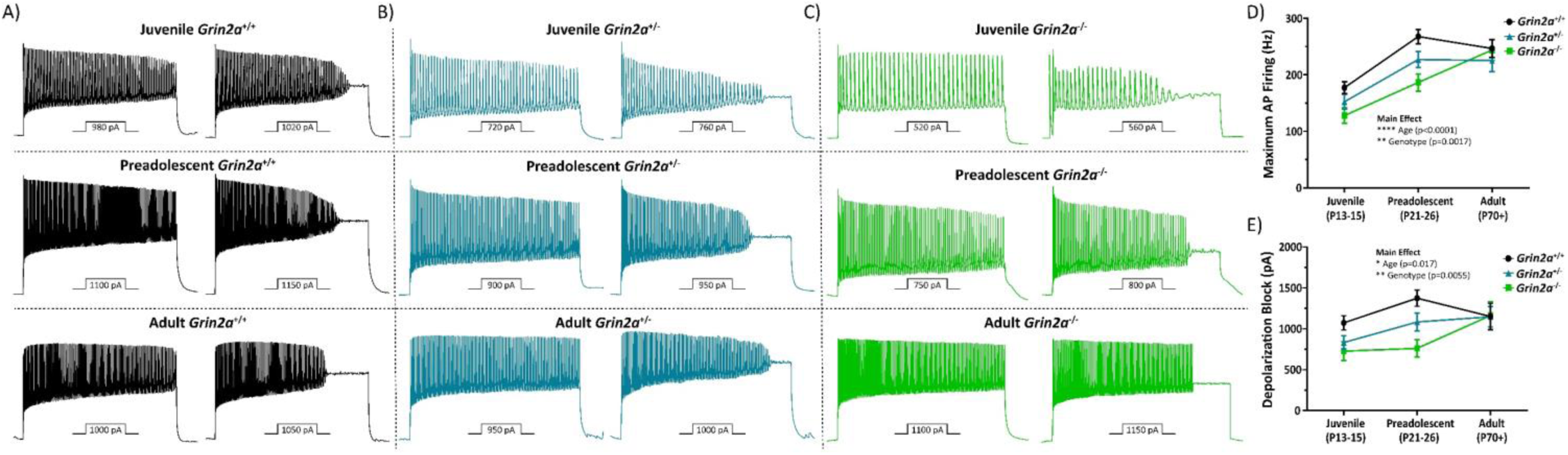
The loss of *Grin2a* causes a transient change in action-potential firing properties of CA1 PV cells. Representative action-potential trains elicited by current injections depicted below each train to illustrate the maximum action-potential firing frequency and the current required for depolarization-induced block of action-potential firing for **A)** *Grin2a^+/+^*, **B)** *Grin2a^+/-^*, and **C)** *Grin2a^-/-^* mice during development. **D)** Maximum action potential firing frequencies show a significant main effect for both age (F_2, 135_ = 29.8; p<0.0001; two-way ANOVA) and genotype (F_2, 135_ = 6.7; p=0.0017; two-way ANOVA). **E)** Currents required to reach depolarization-induced block of action potential firing show a significant main effect for both age (F_2, 135_ = 4.2; p=0.017; two-way ANOVA) and genotype (F_2, 135_ = 5.4; p=0.006; two-way ANOVA). Symbols are mean ± SEM. AP = action-potential; depolarization block = current required to reach depolarization-induced block of action-potential firing; * = p<0.05; ** = p<0.01; ***p<0.001; **** = p<0.0001.

The sum of these suggests that the loss of *Grin2a* impacts the electrophysiological maturation programs of CA1 PV cells, however, this delay is only transient as all perturbed measures eventually attain *Grin2a^+/+^* levels. Immature PV cells may be fast to fire given their increased input resistance, but they are also fast to retire given their alterations in depolarization-induced block and action-potential firing frequency. That is, inhibitory firing will fade in response to high frequency firing, as might occur during initial runup toward seizure initiation. Diminished inhibitory output of a prominent interneuron subtype (PV) could have profound consequences on circuit excitability and network wiring during the critical plasticity period of preadolescent development. These alterations in circuit function could then manifest as epileptiform activity, maladaptive plasticity, or cognitive impairment. Moreover, if the transient nature of these reported alterations in PV cell function also occur in PV interneurons from *GRIN2A* null variant patients, this could provide a molecular mechanism driving hyperexcitability and hypersynchronous epileptiform activity in these patients.

## Discussion

The most important finding of this study is that disease-associated null *GRIN2A* human patients display a largely transient seizure burden that can be resolved with age. To explore this new clinical finding at that circuit and cellular level, we conducted studies using *Grin2a^+/-^* and *Grin2a^-/-^*mice at various stages during neurodevelopment. By utilizing the weighted tau of evoked NMDAR-mediated EPSCs from *Grin2a^+/+^*, *Grin2a^+/-^*and *Grin2a^-/-^* mice we determined that the juvenile (P14-15) time window was the earliest neurodevelopmental stage in which loss of GluN2A-mediated signaling would impart increased circuit excitability in terms of overall charge transfer. Here, the loss of GluN2A-containing NMDARs, which have the fastest deactivation kinetics of all four GluN2 subunits (Hansen et al. 2021), will likely increase the excitatory tone onto every neuron that typically expresses GluN2A. Moreover, we show using a multiple stimulus action-potential spiking paradigm, increased circuit excitability and CA1 pyramidal cell output in juvenile mice of both *Grin2a^+/-^* and *Grin2a^-/-^* mice. These alterations in somatic spiking are not due to global upregulation of other *GRIN* genes (including *Grin2b*) nor can they be attributed to perturbations in the intrinsic excitability or action-potential firing properties of CA1 pyramidal cells.

Deeper evaluation of the developing CA1 circuit lead us to uncover age- and *Grin2a* gene dosing-dependent transient delays in the electrophysiological maturation programs of PV interneurons. Overall, we report that *Grin2a^+/+^* mice reach electrophysiological maturation between the neonatal and juvenile neurodevelopmental timepoints (except for maximum action-potential firing frequency), with *Grin2a^+/-^*mice not reaching electrophysiological maturation until preadolescence, and *Grin2a^-/-^* not reaching electrophysiological maturation until adulthood. The significance of these findings are multifaceted, with the most important being that these data may represent a molecular mechanism describing the transient nature of disease-associated null *GRIN2A* patients. Many questions remain unexplored, however, including how this transient delay is overcome purely with aging. Moreover, other aspects of circuit function are still likely perturbed despite PV cells from *Grin2a^+/-^* and *Grin2a^-/-^*mice attaining *Grin2a^+/+^* electrophysiological function in a delayed fashioned.

Additionally, the data we report here suggest multiple potential roles for the GluN2A subunit of NMDARs in the regulation of function, development, and maturation of GABAergic interneurons in the mouse hippocampus. Specifically, we report that full loss of the *Grin2a* causes aberrations in PV cell density. The loss of *Grin2a* only impacts PV cell density, but not CCK cell density, which may suggest that MGE-derived interneurons are preferentially affected. Our data suggest that the loss of GluN2A may be impacting interneuron survival or apoptosis which are controlled by a combination of internal genetic cues, external neurotropic factors, and early interneuron electrical activity (Southwell et al. 2012; Denaxa et al. 2018; Priya et al. 2018). However, the extent to which local electrical signaling, especially from NMDARs, within PV interneurons controls pro-apoptotic pathways has not been determined. The lack of effect on CCK cell density in *Grin2a*^-/-^ mice could be due to CCK cells having little functional GluN2A expression (Perszyk et al. 2016; Booker et al. 2021; Matta et al. 2013). In addition to changes in PV cell density, we also report that the total loss of *Grin2a* generates an upregulation of *Grin2d* and *Grin3a* mRNA. Given that PV cells highly express GluN2D-containing NMDARs (Garst-Orozco et al. 2020; Hanson et al. 2019), these two findings may coincide. However, there is little evidence of *Grin3a* expression on PV cells (Murillo et al. 2021; Bossi et al. 2022). Instead, recent studies highlight immense GluN3A-mediated NMDA currents on somatostatin interneurons (Bossi et al. 2022), which are also generated from the MGE. Thus, PV cells may not be the only interneuron subtype that is dysfunctional in *Grin2a^-/-^* mice. Future experiments should focus on elucidating electrophysiological function of other MGE-derived interneurons, such as somatostatin cells and some neurogliaform cells.

We also show that CA1 PV cells from *Grin2a*^+/+^ mice display age-dependent alterations in passive electrical and action-potential firing properties. By examining both passive intrinsic properties and action potential firing characteristics, we can estimate how these cells may respond to excitatory afferent activity and how efficiently they can transduce somatic depolarizations into signal-carrying action potentials. In both *Grin2a^+/-^* and *Grin2a^-/-^*mice we show age- and *Grin2a* gene dosing-dependent transient prolongations of the membrane time constant, increases in input resistance, and longer action potential half-widths. Each of these electrophysiological measures are known to be age-dependent, with PV cells in neonatal mice displaying a higher input resistance, longer membrane time constant, and longer action potential half-widths than PV cells from preadolescent mice (Okaty et al. 2009; Miyamae et al. 2017; Doischer et al. 2008; Goldberg et al. 2011). Moreover, PV cells from *Grin2a^+/-^* and *Grin2a*^-/-^ mice have a lower thresholds to reach depolarization-induced block of action potential firing and lower peak action potential firing frequencies. These latter electrophysiological characteristics could lead to failure of circuit level inhibition as interneuron depolarization increases, consistent with a hyperexcitable phenotype.

The exact mechanism for this transcriptional shift from immature to mature PV electrophysiological properties is not fully understood but has been shown to be activity-dependent (Miller et al. 2011; Dehorter et al. 2015). Thus, the loss of GluN2A-mediated signaling appears to slow cellular maturation of CA1 PV cells. However, it is currently unknown whether these delays stem from a loss of GluN2A-mediated signaling on PV cell dendrites (cell autonomous) or due to excitability changes of local pyramidal cells (non-cell autonomous). All measure electrophysiological properties ultimately reach *Grin2a^+/+^*levels in adult mice, suggesting that although GluN2A signaling may be sufficient to initiate these maturation transcriptional programs, it is not required. While the overall effects of a prolonged immature electrophysiological profile in developing PV cells are not known, this period coincides with critical period plasticity in various brain regions (Hensch 2005; Takesian and Hensch 2013), and thus, could critically alter circuit formation in the hippocampus, and other regions. Specifically, larger input resistances and lower peak action potential firing frequencies will likely dampen the temporal resolution of inhibitory tone in a developing network, which may promote maladaptive plasticity and incorrect circuit wiring. Additionally, changes in depolarization-induced block of action potentials correlates with neural excitability and propagation of epileptiform activity (Calin, Ilie, and Akerman 2021). Thus, disruption of PV cell function during development could have profound consequences for eventual mature networks that arise.

## Acknowledgements

Research reported in this publication was supported in part by an Emory University Synergy grant awarded to S.F.T. and Children’s Healthcare of Atlanta as well as by the Emory University Integrated Cellular Imaging Core and the Emory Integrated Genomics Core, which are subsidized by the Emory University School of Medicine and part of the Emory Integrated Core Facilities. This work was supported by the following grants from the National Institutes of Health, National Institute of Neurological Disease and Stroke: NS113530 (C.R.C.), MH127404 (H.Y.), HD082373 (H.Y.), and NS111619 (S.F.T.). T.A.B was supported by the Ponzio Family Chair in Neurology Research from the Children’s Hospital Colorado Foundation. K.L.P. and T.A.B. were also supported by Simon’s Foundation. I.H. was supported by The Hartwell Foundation (Individual Biomedical Research Award), NINDS (K02NS112600, U24NS120854-01, U54NS108874-04), the Eunice Kennedy Shriver National Institute of Child Health and Human Development through the Children’s Hospital of Philadelphia and the University of Pennsylvania (U54HD086984), the German Research Foundation (HE5415/3-1, HE5415/5-1, HE5415/6-1, HE5415/7-1), the National Center for Advancing Translational Sciences of the NIH (UL1TR001878), the Institute for Translational Medicine and Therapeutics’ (ITMAT) at the Perelman School of Medicine of the University of Pennsylvania, and by Children’s Hospital of Philadelphia through the Epilepsy NeuroGenetics Initiative (ENGIN). The content is solely the responsibility of the authors and does not necessarily reflect the official views of the National Institute of Health. The authors would also like to thank Steven Hunt and Andrew Hancock for their assistance with interneuron morphological analysis, and Nicholas Varvel and Asheebo Rojas for their assistance with general immunohistochemistry. Additionally, the authors would like to thank Masayoshi Mishina and Stefano Vicini for generously sharing the *Grin2a*^-/-^ mice, and Jing Zhang for excellent technical assistance.

---

**Supplemental Figure S1.**
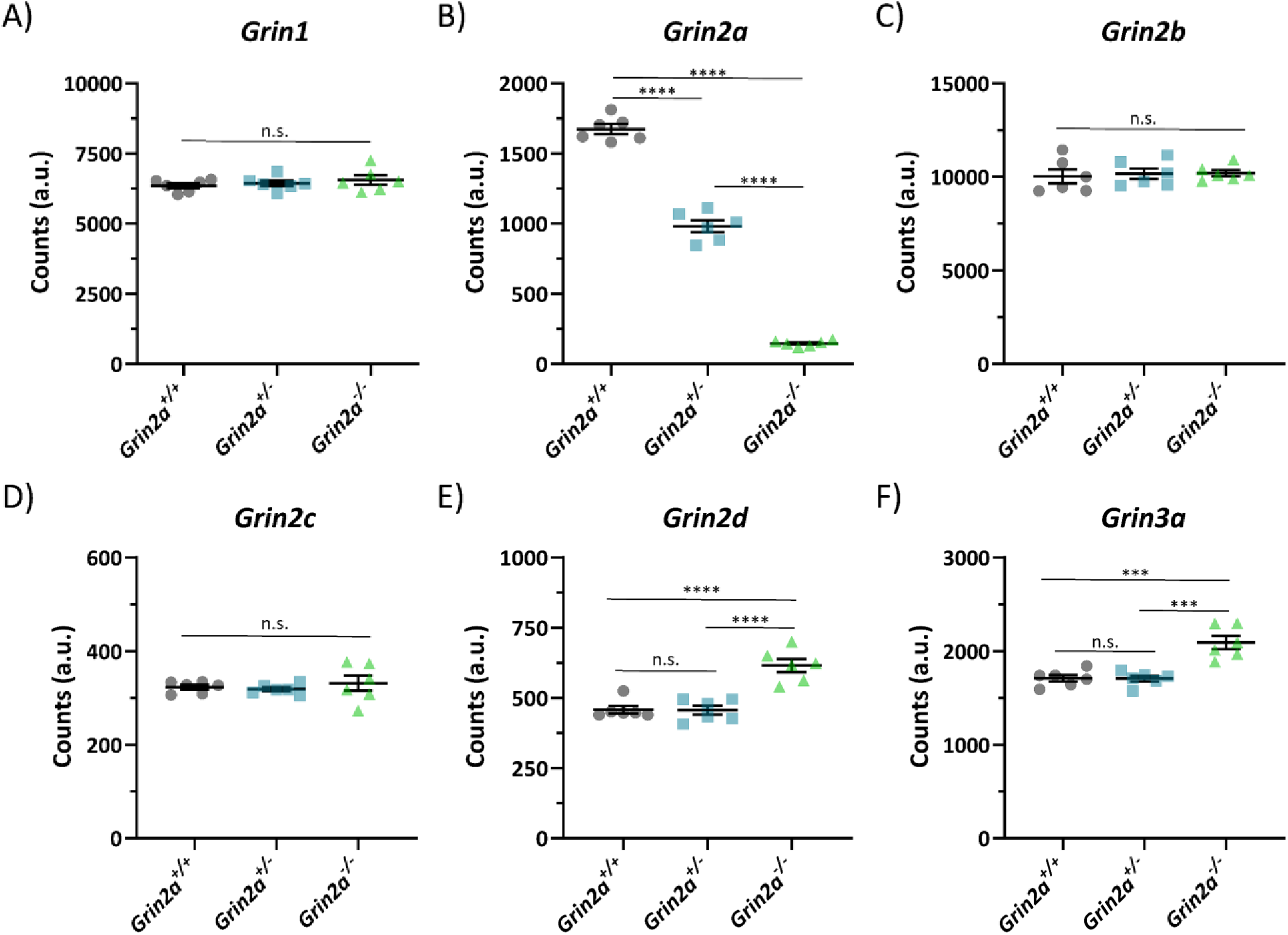
Evaluation of all *GRIN* gene transcripts in developing hippocampus. Whole hippocampi from juvenile mice across 18 samples, six from each genotype, were subjected to NanoString gene expression analysis. Normalized raw counts were then plotted and tested for statistical significance, controlling for false-discovery rate. There were no significant differences in **A)** *Grin1*, **C)** *Grin2b*, and **D)** *Grin2c* gene expression across all three genotypes. As expected, **B)** *Grin2a* gene expression shows a statistically significant, genotype-dependent decrease. Both **E)** *Grin2d* and **F)** *Grin3a* genes were statistically significantly upregulated in *Grin2a^-/-^* mice compared to *Grin2a^+/+^*, with no significant differences between *Grin2a^+/+^* and *Grin2a^+/-^* mice. Counts for the *Grin3b* gene fell below our limit of detection as determined by negative controls and are not shown. All statistical tests performed were one-way ANOVAs, with multiple post-hoc comparisons. *** = p<0.001; **** = p<0.0001; n.s. = not significant; a.u. = arbitrary units.

**Supplemental Figure S2.**
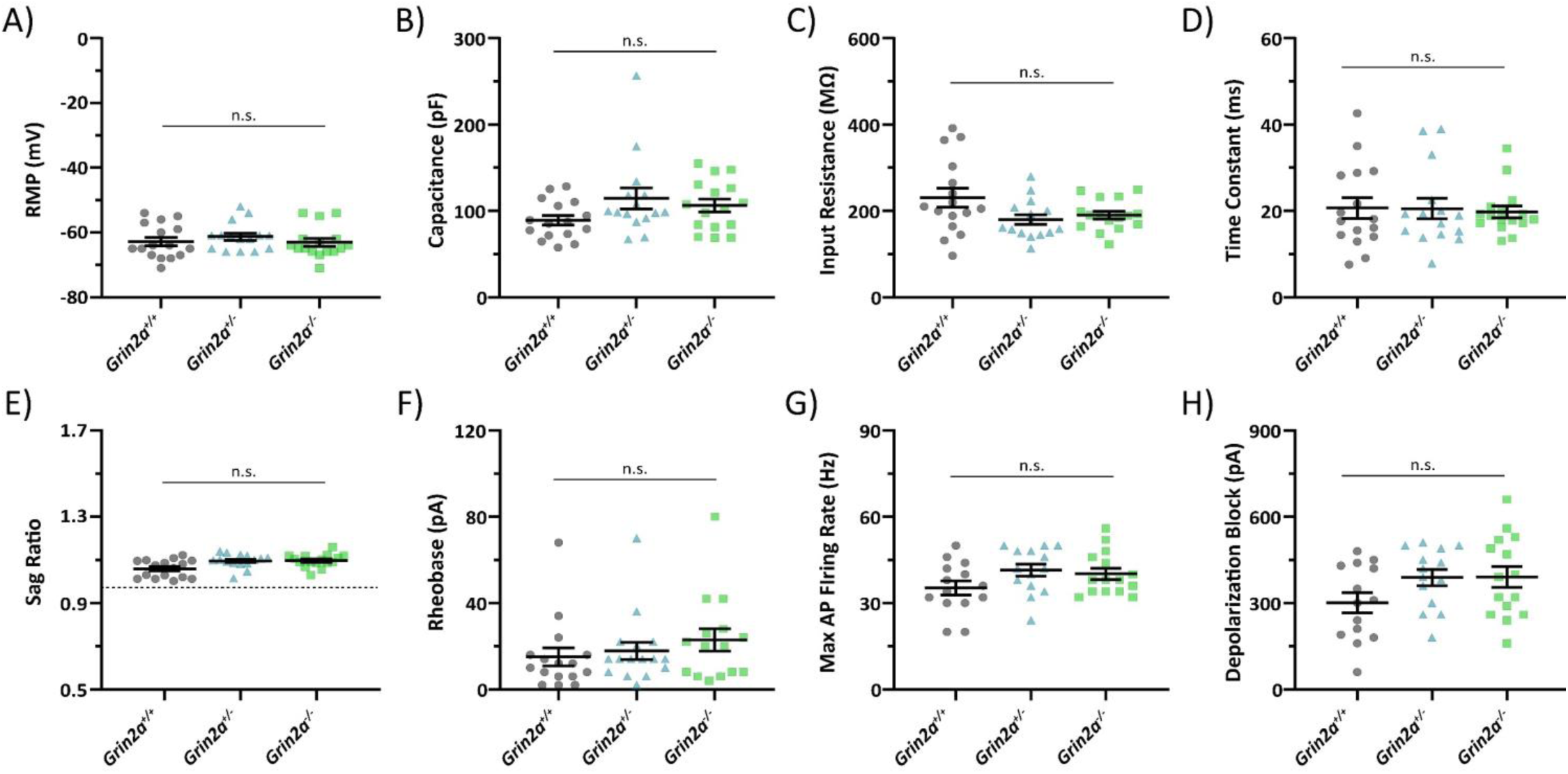
Juvenile CA1 circuit hyperexcitability is not due to changes in CA1 pyramidal cell intrinsic or action-potential firing properties. Data from juvenile (P14-16) CA1 pyramidal cells show no significant differences in **A)** resting membrane potential, **B)** cell capacitance, **C)** input resistance, **D)** membrane time constant, **E)** sag ratio, **F)** rheobase, **G)** maximum action-potential firing rate, or **H)** current required to reach depolarization-induced blockade of action-potential firing when assayed via one-way ANOVA. Data represented show mean ± SEM. RMP = resting membrane potential; AP = action-potential; depolarization block = current required to reach depolarization-induced blockade of action-potential firing; n.s. = not significant.

**Supplemental Figure S3.**
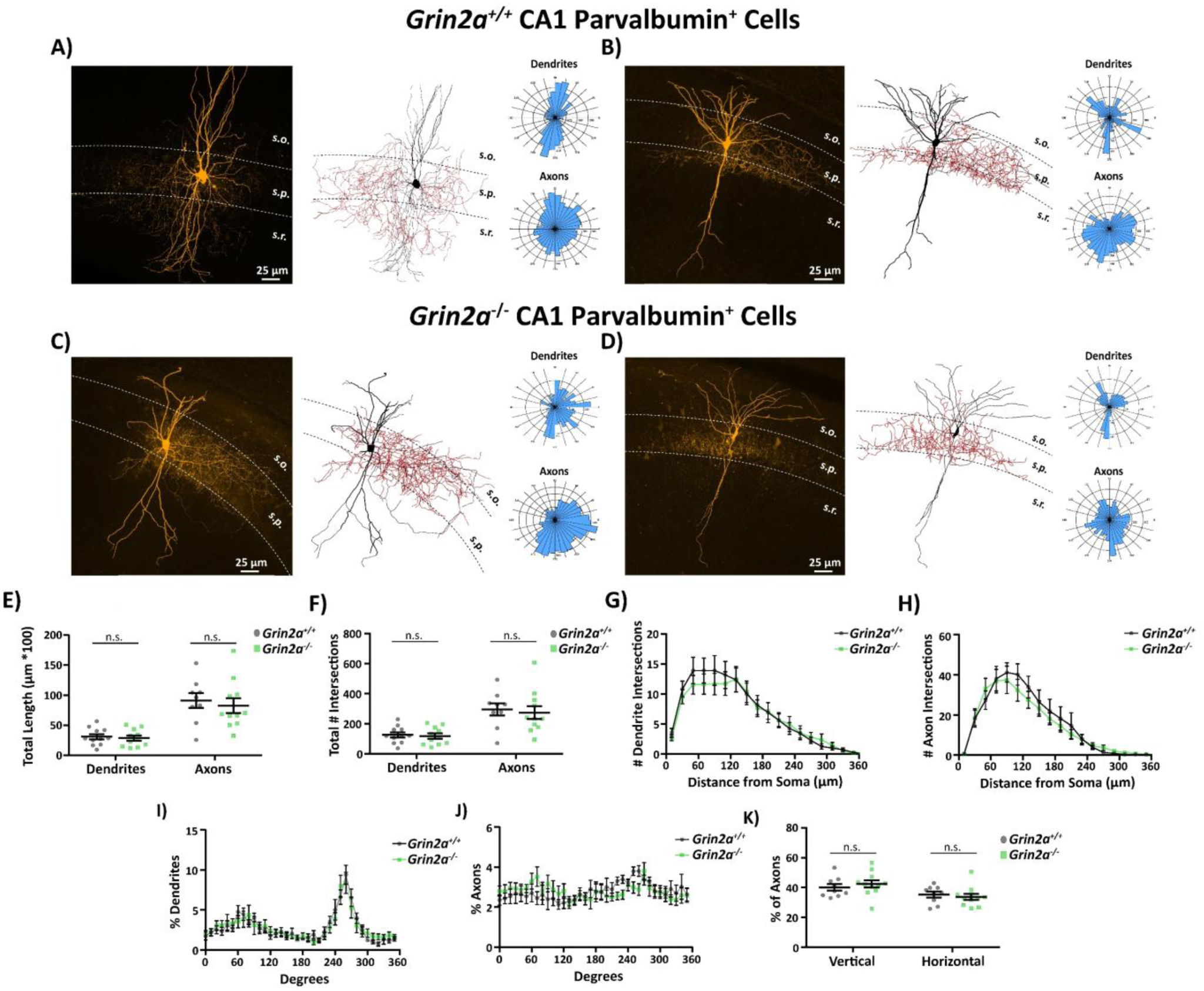
There is no difference in CA1 PV cell morphology in preadolescent *Grin2a*^-/-^ mice. Representative images for **A-B)** *Grin2a^+/+^*or **C-D)** *Grin2a*^-/-^ biocytin backfilled CA1 PV cells shown as maximum intensity projection images with accompanying NeuroLucida tracings (black for dendrites and red for axons) and polar plots for both dendrites and axons. There is no change in **E)** total length of dendrites or axons or in **F)** the total number of Sholl intersections in dendrites or axons in preadolescent CA1 PV cells from either genotype. Sholl analysis of both **G)** dendrites and **H)** axons from preadolescent CA1 PV cells are also unchanged across genotype. The placement of both **I)** dendrites or **J)** axons assessed via polar plots showed no differences, with **K)** equal axonal placement in both the vertical and horizontal planes. Data obtained from 9-11 slices across 4 animals per genotype. Symbols are mean ± SEM. n.s. = not significant. s.o. = *stratum oriens*; s.p. = *stratum pyramidale*; s.r. = *stratum radiatum*

**Supplemental Figure S4.**
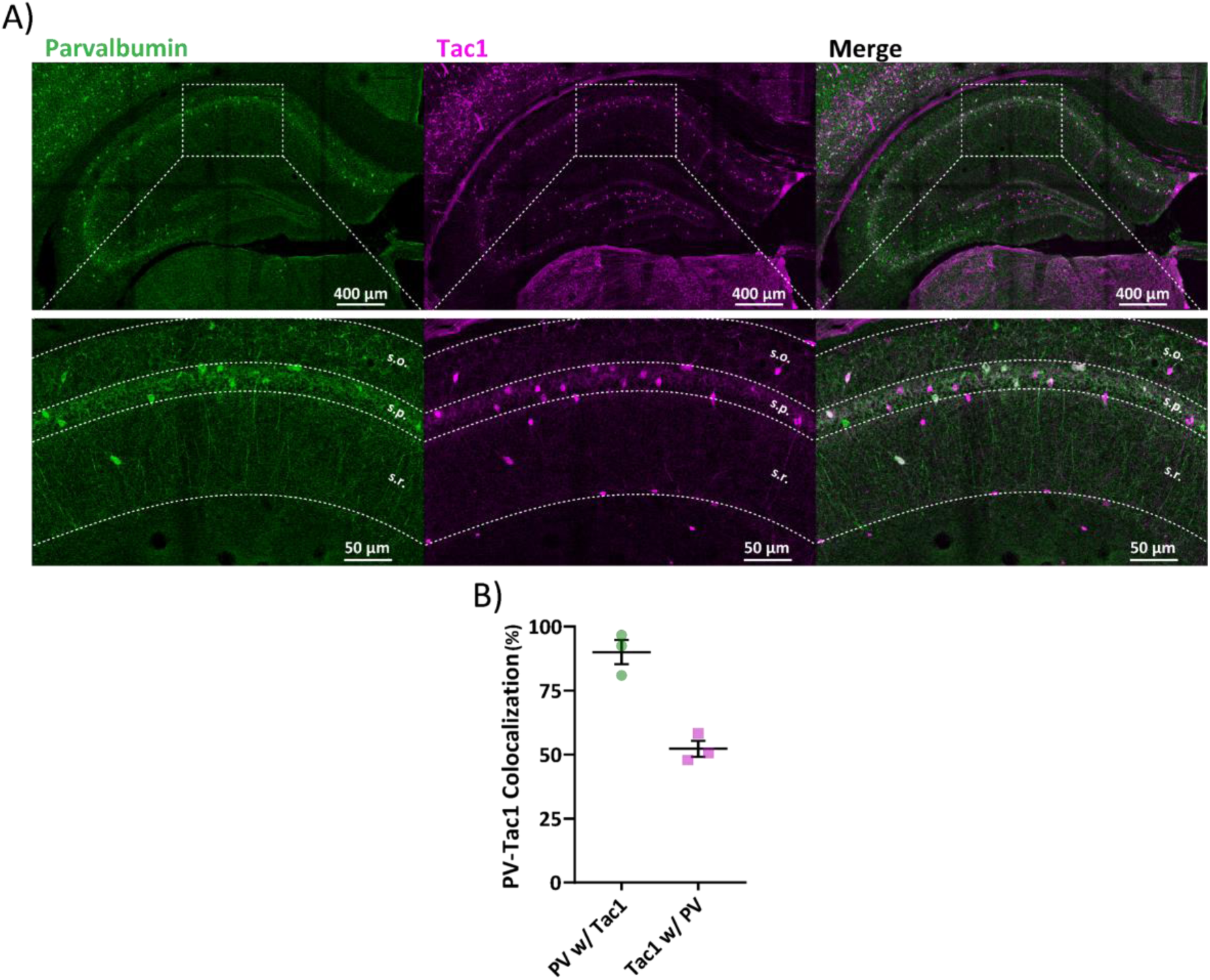
The majority of cells that are immunopositive for parvalbumin are also immunopositive for Tac1. **A)** Example hippocampal slice from an adult *Tac1*-Cre × Floxed-eGFP mouse that has been stained for anti-parvalbumin and anti-eGFP. Inset displays a more detailed view of CA1, with the majority of PV-positive cells in *stratum oriens* and *stratum pyramidale* also being immunopositive for Tac1. **B)** Quantification of PV and Tac1 co-expression overlap. Since CA1 *stratum oriens* and *stratum pyramidale* had the most extensive overlap for PV and Tac1, all recordings made from *Tac1*-positive cells were chosen in these two layers. *s.o.* = *stratum oriens*; *s.p.* = *stratum pyramidale*; *s.r.* = *stratum radiatum*.

**Supplemental Figure S5.**
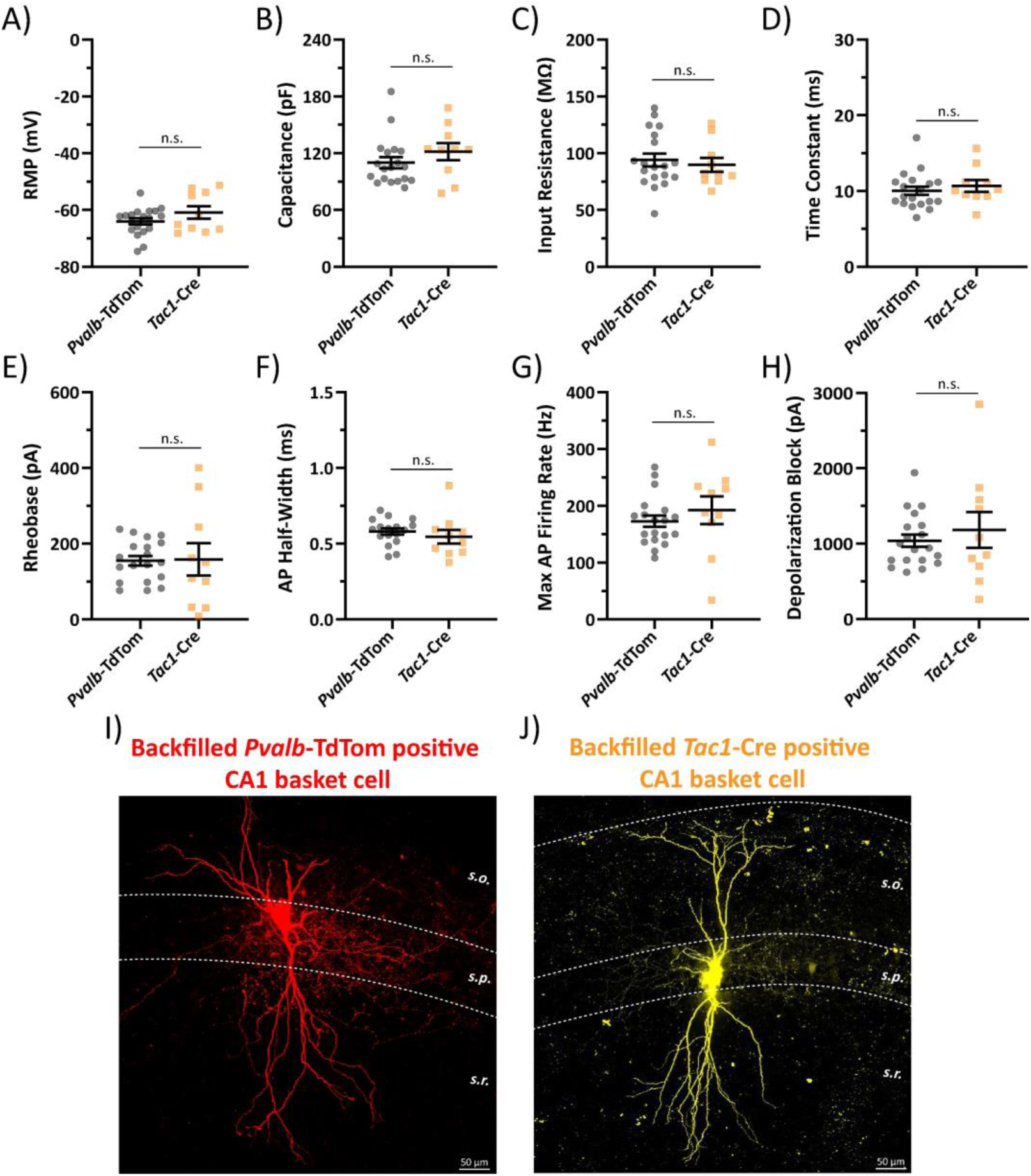
*Pvalb*-TdTom identified CA1 PV interneurons display identical electrophysiological properties as CA1 PV interneurons identified via *Tac1*-Cre. **A-H)** Both passive and action-potential firing electrophysiological properties of juvenile CA1 PV cells from *Pvalb*-TdTom and *Tac1*-Cre show no statistically significant differences. Example backfills of CA1 PV cells using **I)** *Pvalb*- TdTom and **J)** *Tac1*-Cre mouse lines indicates that both can be reliably used to visualize PV basket cells in CA1 *stratum pyramidale*. RMP = resting membrane potential; AP = action-potential; n.s. = not significant; *s.o.* = *stratum oriens*; *s.p.* = *stratum pyramidale*; *s.r.* = *stratum radiatum*.

**Supplemental Table S1.**
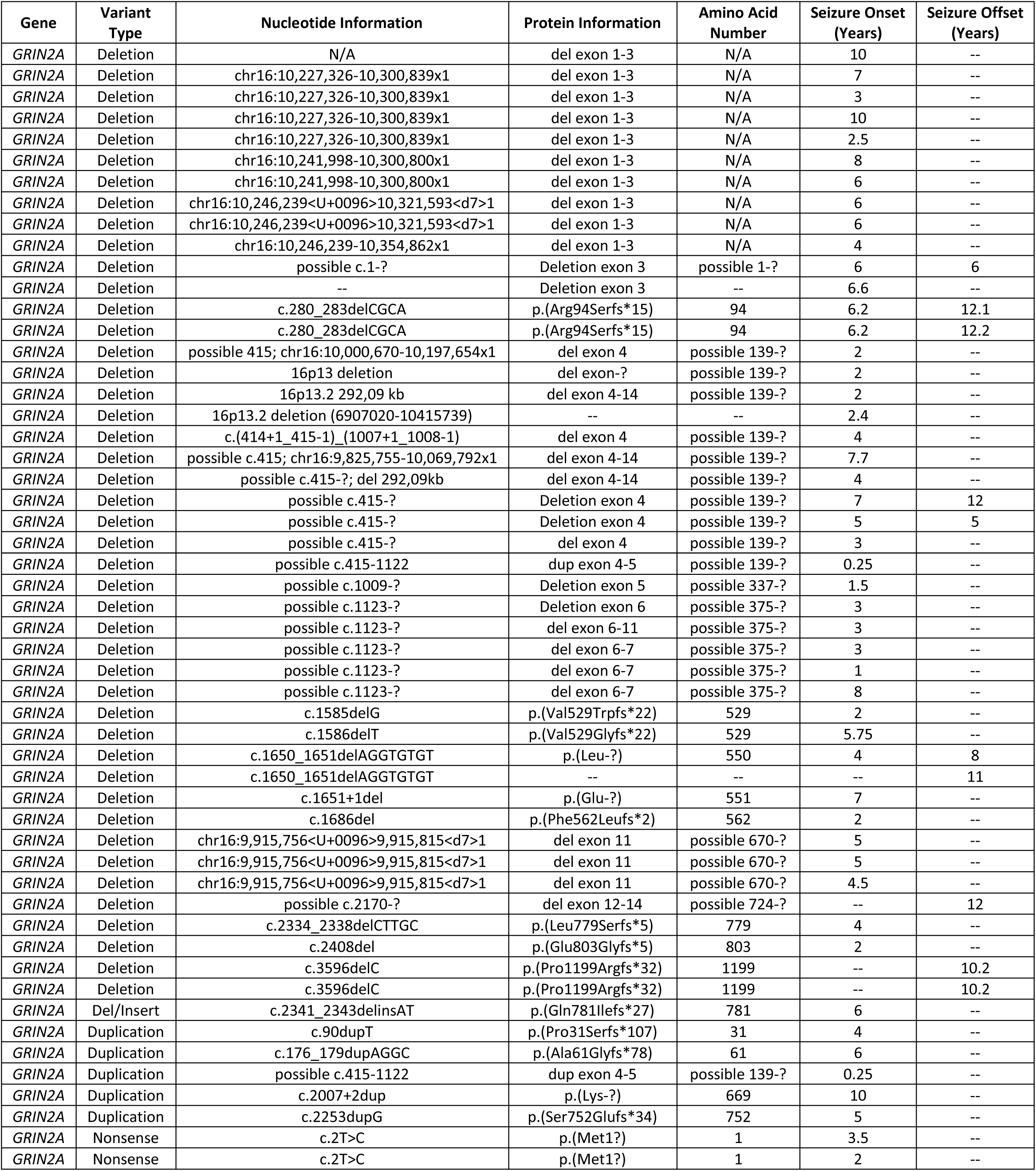
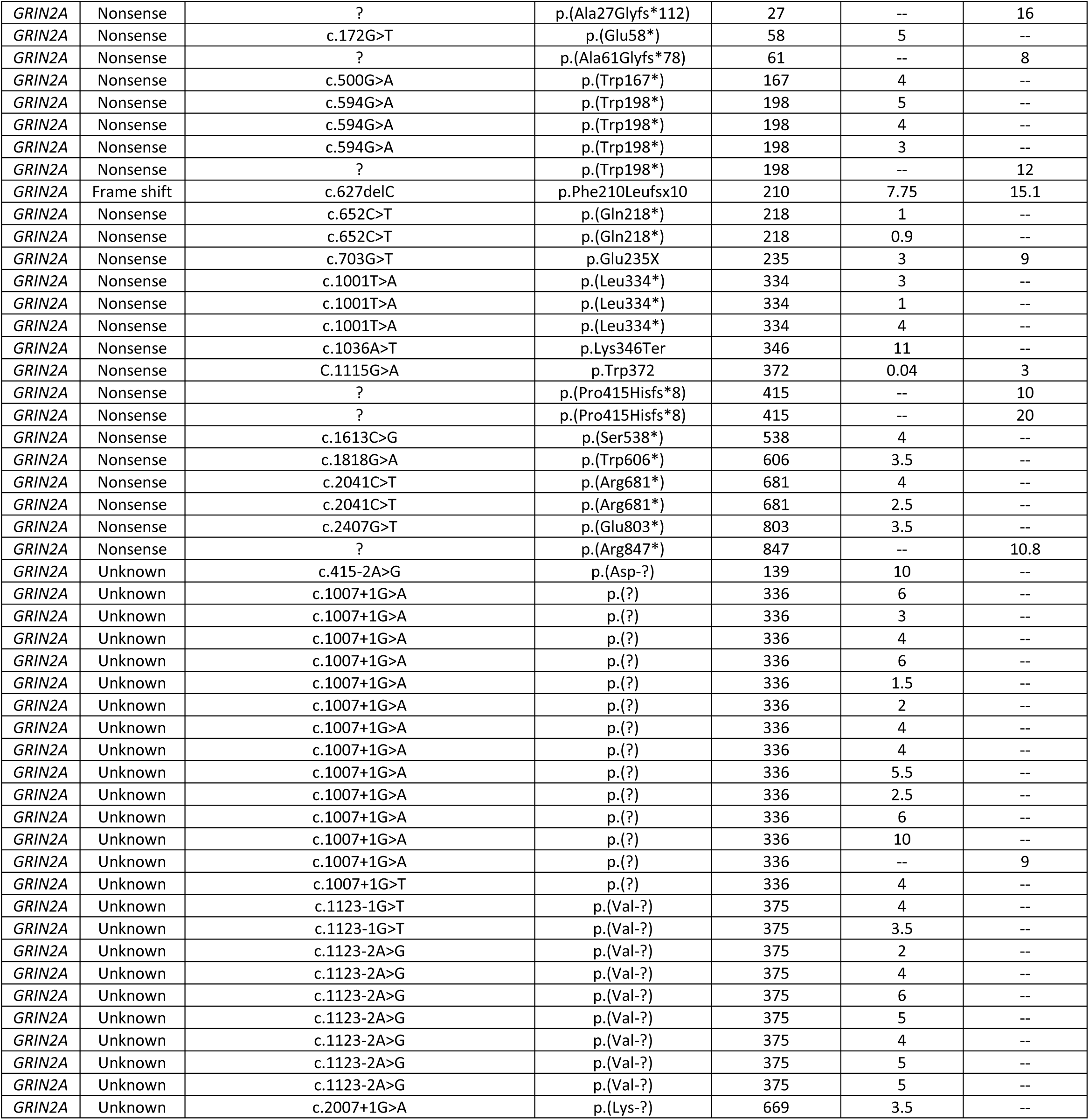
Null *GRIN2A* variant patient data with time when seizures began and/or ended. Data represented are gathered from the *GRIN* portal (https://grin-portal.broadinstitute.org/), or respective patient registries housed at University of Leipzig, Children’s Hospital of Philadelphia, or University of Colorado School of Medicine. Seizure offset is defined as freedom from seizures for two consecutive years. N/A = not applicable. ‘--’ = data not available at time of publication. Some patients had not reached an age where offset was expected.

**Supplemental Table S2.** Missense *GRIN2A* variant patient data with information on when seizure burden began and/or ended. Data represented are gathered from the *GRIN* portal (https://grin-portal.broadinstitute.org/), University of Leipzig, Children’s Hospital of Philadelphia, or University of Colorado Anshutz Medical Center. Seizure offset is defined as freedom from seizures for two consecutive years. ‘--’ = data not available at time of publication.

| Gene | Variant Type | Nucleotide Information | Protein Information | Amino Acid Number | Seizure Onset (Years) | Seizure Offset (Years) |
| --- | --- | --- | --- | --- | --- | --- |
| *GRIN2A* | Missense | c.1667C>T | p.(Arg217Trp) | 217 | 0.25 | 1.7 |
| *GRIN2A* | Missense | c.692G>A | p.(Cys231Tyr) | 231 | 3 | -- |
| *GRIN2A* | Missense | c.1232T>A | p.(Leu411Gln) | 411 | 3 | -- |
| *GRIN2A* | Missense | c.1251C>T | p.(Val417Val) | 417 | 0.8 | -- |
| *GRIN2A* | Missense | c.1306T>C | p.(Cys436Arg) | 436 | 4 | -- |
| *GRIN2A* | Missense | c.1447G>A | p.(Gly483Arg) | 483 | 4 | -- |
| *GRIN2A* | Missense | c.1492G>A | p.(Gly498Ser) | 498 | 8 | -- |
| *GRIN2A* | Missense | c.1532C>T | p.(Ser511Leu) | 511 | 2.5 | -- |
| *GRIN2A* | Missense | c.1552C>T | p.(Arg518Cys) | 518 | 3 | -- |
| *GRIN2A* | Missense | c.1553G>A | p.(Arg518His) | 518 | 3 | -- |
| *GRIN2A* | Missense | c.1553G>A | p.(Arg518His) | 518 | 2 | -- |
| *GRIN2A* | Missense | c.1592C>T | p.(Thr531Met) | 531 | 6.7 | -- |
| *GRIN2A* | Missense | c.1592C>T | p.(Thr531Met) | 531 | 7 | -- |
| *GRIN2A* | Missense | c.1592C>T | p.(Thr531Met) | 531 | 11 | -- |
| *GRIN2A* | Missense | c.1642G>A | p.(Ala548Thr) | 548 | 6 | -- |
| *GRIN2A* | Missense | c.1655C>G | p.(Pro552Arg) | 552 | 0.75 | -- |
| *GRIN2A* | Missense | c.1655C>G | p.(Pro552Arg) | 552 | 0.75 | -- |
| *GRIN2A* | Missense | c.1841A>G | p.(Asn614Ser) | 614 | 0.4 | -- |
| *GRIN2A* | Missense | c.1845C>A | p.(Asn615Lys) | 615 | 0.75 | -- |
| *GRIN2A* | Missense | c.1845C>A | p.(Asn615Lys) | 615 | 0.25 | -- |
| *GRIN2A* | Missense | c.1903G>A | p.(Ala635Thr) | 635 | 0.6 | -- |
| *GRIN2A* | Missense | c.1930A>G | p.(Ser644Gly) | 644 | 0.5 | -- |
| *GRIN2A* | Missense | c.1936A>G | p.(Thr646Ala) | 646 | 0.7 | -- |
| *GRIN2A* | Missense | c.1936A>G | p.(Thr646Ala) | 646 | 0.25 | -- |
| *GRIN2A* | Missense | c.1945C>G | p.(Leu649Val) | 649 | 0.75 | -- |
| *GRIN2A* | Missense | c.1946_1947delin | p.(Leu649Pro) | 649 | 1 | -- |
| *GRIN2A* | Missense | c.1948G>T | p.(Ala650Ser) | 650 | 1.5 | -- |
| *GRIN2A* | Missense | c.1954T>G | p.(Phe652Val) | 652 | 2 | -- |
| *GRIN2A* | Missense | c.1961T>C | p.(Ile654Thr) | 654 | 0.2 | -- |
| *GRIN2A* | Missense | c.2081T>C | p.(Ile694Thr) | 694 | 2.5 | -- |
| *GRIN2A* | Missense | c.2095C>T | p.(Pro699Ser) | 699 | 8 | -- |
| *GRIN2A* | Missense | c.2113A>G | p.(Met705Val) | 705 | 8 | -- |
| *GRIN2A* | Missense | c.2191 G>A | p.Asp731Asn | 731 | 1.9 | -- |
| *GRIN2A* | Missense | c.2191G>A | p.(Asp731Asn) | 731 | 4 | -- |
| *GRIN2A* | Missense | c.2191G>A | p.Asp731Asn | 731 | 3.9 | -- |
| *GRIN2A* | Missense | c.2326G>T | p.(Asp776Tyr) | 776 | 7 | -- |
| *GRIN2A* | Missense | c.2434C>A | p.(Leu812Met) | 812 | 1.5 | -- |
| *GRIN2A* | Missense | c.2449A>G | p.(Met817Val) | 817 | 2 | -- |
| *GRIN2A* | Missense | c.2450T>C | p.(Met817Thr) | 817 | 2 | -- |
| *GRIN2A* | Missense | c.2450T>C | p.(Met817Thr) | 817 | 1.25 | -- |
| *GRIN2A* | Missense | c.2452G>A | p.(Ala818Thr) | 818 | 3 | -- |
| *GRIN2A* | Missense | c.2453C>A | p.(Ala818Glu) | 818 | 1 | -- |
| *GRIN2A* | Missense | c.2907C>G | p.(Asn969Lys) | 969 | 2.5 | -- |
| *GRIN2A* | Missense | c.3664C>G | p.(Pro1222Ala) | 1222 | 7.9 | -- |
| *GRIN2A* | Missense | c.3664C>G | p.(Pro1222Ala) | 1222 | 7.8 | -- |

**Supplemental Table S3.** Action-potential spiking probability in *Grin2a^+/+^*, *Grin2a^+/-^*, and *Grin2a^-/-^* juvenile (P14-16) mice. Data represented are mean ± SEM. AP = action potential.

|  | AP Probability<br>1<sup>st</sup> Stimulation | AP Probability<br>2<sup>nd</sup> Stimulation | AP Probability<br>3<sup>rd</sup> Stimulation | AP Probability<br>4<sup>th</sup> Stimulation | AP Probability<br>5<sup>th</sup> Stimulation | Total Number of<br>APs Generated |
| --- | --- | --- | --- | --- | --- | --- |
| *Grin2a*<sup>+/+</sup> mice<br>N=16 cells/5 mice | 0.05 ± 0.04 | 0.17 ± 0.07 | 0.2 ± 0.07 | 0.08 ± 0.03 | 0.05 ± 0.05 | 2.75 ± 0.91 |
| *Grin2a*<sup>+/-</sup> mice<br>N=16 cells/4 mice | 0.06 ± 0.02 | 0.58 ± 0.1 | 0.29 ± 0.09 | 0.13 ± 0.05 | 0.1 ± 0.04 | 5.88 ± 1.18 |
| *Grin2a*<sup>-/-</sup> mice<br>N=16 cells/4 mice | 0.03 ± 0.04 | 0.51 ± 0.1 | 0.38 ± 0.09 | 0.21 ± 0.06 | 0.09 ± 0.04 | 6.06 ± 1.12 |

**Supplemental Table S4.** Passive and action-potential firing properties of CA1 pyramidal cells in *Grin2a^+/+^*, *Grin2a^+/-^*, and *Grin2a^-/-^* juvenile (P14-16) mice. Data represented are mean ± SEM. RMP = resting membrane potential; AP = action potential; depolarization block = current required for depolarization-induced block of AP firing.

|  | RMP<br>(mV) | Capacitance<br>(pF) | Input Resistance<br>(MΩ) | Time Constant<br>(ms) | Rheobase<br>(pA) | Sag Ratio | Max AP Firing<br>(Hz) | Depolarization Block<br>(pA) |
| --- | --- | --- | --- | --- | --- | --- | --- | --- |
| *Grin2a*<sup>+/+</sup> mice<br>N=16 cells/5 mice | -62 ± 1.3 | 89 ± 5.8 | 231 ± 22 | 21 ± 2.4 | 20 ± 4.4 | 1.06 ± 0.01 | 35 ± 2.4 | 301 ± 35 |
| *Grin2a*<sup>+/-</sup> mice<br>N=15 cells/4 mice | -61 ± 1.1 | 114 ± 12 | 179 ± 12 | 21 ± 2.4 | 23 ± 5.2 | 1.1 ± 0.01 | 41 ± 2 | 389 ± 27 |
| *Grin2a*<sup>-/-</sup> mice<br>N=16 cells/4 mice | -63 ± 1.2 | 106 ± 7.3 | 190 ± 8.9 | 20 ± 1.4 | 19 ± 4 | 1.1 ± 0.01 | 40 ± 1.9 | 391 ± 36 |

**Supplemental Table S5.** Parvalbumin-positive CA1 density and cellular lamination in *Grin2a^+/+^*, *Grin2a^+/-^*, and *Grin2a^-/-^* preadolescent (P21-26) mice. Data represented are mean ± SEM. 4-5 hippocampi from each animal were imaged across 4 animals per genotype for all staining and cell counts.

|  | CA1 Cell Density<br>(# / mm<sup>3</sup>) | *stratum oriens*<br>(% total CA1 PV cells) | *stratum pyramidale*<br>(% total CA1 PV cells) | *stratum radiatum*<br>(% total CA1 PV cells) | *stratum lacuosum moleculare*<br>(% total CA1 PV cells) |
| --- | --- | --- | --- | --- | --- |
| *Grin2a*<sup>+/+</sup> mice<br>N=17 slices/4 mice | 4875 ± 162 | 42 ± 2.8 | 51 ± 2.8 | 7.5 ± 1.5 | 0 |
| *Grin2a*<sup>+/-</sup> mice<br>N=16 slices/4 mice | 4542 ± 198 | 40 ± 2.7 | 49 ± 2.4 | 11 ± 8.4 | 0 |
| *Grin2a*<sup>-/-</sup> mice<br>N=20 slices/4 mice | 6488 ± 276 | 40 ± 2.3 | 52 ± 2.3 | 9.4 ± 5.4 | 0 |

**Supplemental Table S6.** Cholecystokinin-positive CA1 density and cellular lamination in *Grin2a^+/+^*, *Grin2a^+/-^*, and *Grin2a^-/-^* preadolescent (P21-26) mice. Data represented are mean ± SEM.

|  | CA1 Cell Density<br>(# / mm<sup>3</sup>) | *stratum oriens*<br>(% total CA1 CCK cells) | *stratum pyramidale*<br>(% total CA1 CCK cells) | *stratum radiatum*<br>(% total CA1 CCK cells) | *stratum lacuosum moleculare*<br>(% total CA1 CCK cells) |
| --- | --- | --- | --- | --- | --- |
| *Grin2a*<sup>+/+</sup> mice<br>N=20 slices/4 mice | 4060 ± 210 | 24 ± 2.4 | 11 ± 2.1 | 47 ± 2.2 | 17 ± 2.4 |
| *Grin2a*<sup>+/-</sup> mice<br>N=16 slices/4 mice | 3790 ± 177 | 23 ± 1.9 | 10 ± 1.7 | 50 ± 1.9 | 17 ± 1.6 |
| *Grin2a*<sup>-/-</sup> mice<br>N=20 slices/4 mice | 4209 ± 299 | 27 ± 2.3 | 16 ± 1.7 | 45 ± 2.8 | 12 ± 9 |

**Supplemental Table S7.**
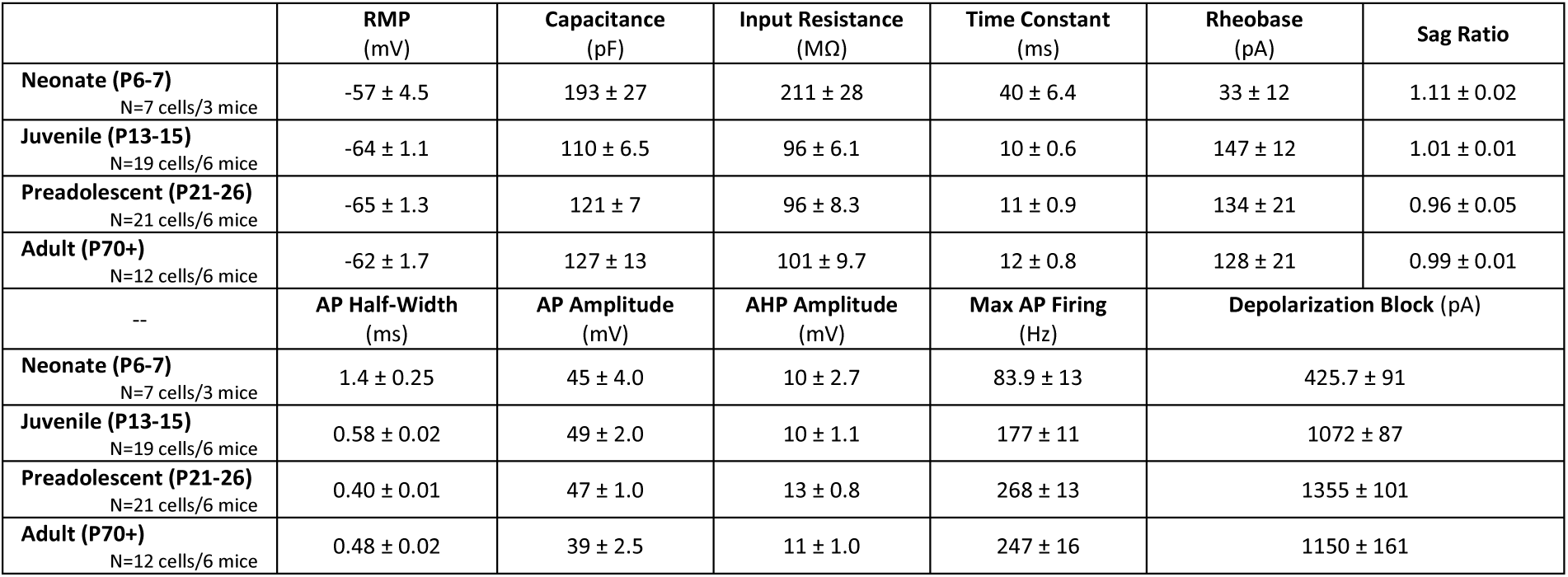
Electrophysiological properties of wildtype CA1 PV cells during development. Data represented are mean ± SEM. RMP = resting membrane potential; AP = action potential; AHP = afterhyperpolarization of the AP; depolarization block = current required for depolarization-induced block of AP firing.

**Supplemental Table S8.** Passive electrophysiological properties during development in *Grin2a^+/+^*, *Grin2a^+/-^*, and *Grin2a^-/-^* CA1 PV cells. RMP = resting membrane potential.

|  | RMP<br>(mV) | Capacitance<br>(pF) | Input Resistance<br>(MΩ) | Time Constant<br>(ms) | Sag Ratio |
| --- | --- | --- | --- | --- | --- |
| Juvenile *Grin2a*<sup>+/+</sup><br>N=19 cells/6 mice | -64 ± 1.1 | 110 ± 6.5 | 96 ± 6.1 | 10 ± 0.6 | 1.01 ± 0.01 |
| Juvenile *Grin2a*<sup>+/-</sup><br>N=19 cells/6 mice | -63 ± 1.6 | 145 ± 8.1 | 133 ± 13 | 19 ± 2.5 | 1.02 ± 0.01 |
| Juvenile *Grin2a*<sup>-/-</sup><br>N=14 cells/4 mice | -59 ± 3 | 111 ± 11 | 184 ± 23 | 22 ± 4.4 | 1.04 ± 0.02 |
| Preadolescent *Grin2a*<sup>+/+</sup><br>N=21 cells/6 mice | -65 ± 1.3 | 121 ± 7 | 96 ± 8.3 | 11 ± 0.9 | 0.96 ± 0.05 |
| Preadolescent *Grin2a*<sup>+/-</sup><br>N=19 cells/5 mice | -61 ± 1.1 | 126 ± 12 | 113 ± 9.4 | 13 ± 1.1 | 1.01 ± 0.004 |
| Preadolescent *Grin2a*<sup>-/-</sup><br>N=19 cells/5 mice | -63 ± 3.8 | 117 ± 10 | 149 ± 17 | 17 ± 2 | 1.01 ± 0.05 |
| Adult *Grin2a*<sup>+/+</sup><br>N=12 cells/6 mice | -62 ± 1.7 | 127 ± 13 | 101 ± 9.7 | 12 ± 0.8 | 0.99 ± 0.01 |
| Adult *Grin2a*<sup>+/-</sup><br>N=12 cells/4 mice | -59 ± 2.5 | 107 ± 5.8 | 99 ± 7.9 | 11 ± 1.0 | 1.01 ± 0.003 |
| Adult *Grin2a*<sup>-/-</sup><br>N=14 cells/5 mice | -62 ± 1.8 | 117 ± 9.3 | 107 ± 12 | 12 ± 0.8 | 1.00 ± 0.002 |

**Supplemental Table S9.** Action-potential waveform and firing properties during development in *Grin2a^+/+^*, *Grin2a^+/-^*, and *Grin2a^-/-^* CA1 PV cells. AP = action potential; AHP = afterhyperpolarization of the AP; depolarization block = current required for depolarization-induced block of AP firing.

|  | Rheobase<br>(pA) | AP Half-Width<br>(mV) | AP Amplitude<br>(mV) | AHP Amplitude<br>(mV) | Max AP Firing<br>(Hz) | Depolarization<br>Block (pA) |
| --- | --- | --- | --- | --- | --- | --- |
| Juvenile *Grin2a*<sup>+/+</sup><br>N=19 cells/6 mice | 147 ± 12 | 0.58 ± 0.02 | 49 ± 2.0 | 10 ± 1.1 | 177 ± 11 | 1072 ± 87 |
| Juvenile *Grin2a*<sup>+/-</sup><br>N=19 cells/6 mice | 100 ± 14 | 0.69 ± 0.04 | 47 ± 1.1 | 11 ± 1.1 | 153 ± 12 | 861 ± 78 |
| Juvenile *Grin2a*<sup>-/-</sup><br>N=14 cells/4 mice | 82 ± 22 | 0.82 ± 0.07 | 46 ± 4.5 | 10 ± 1.3 | 135 ± 15 | 725 ± 116 |
| Preadolescent *Grin2a*<sup>+/+</sup><br>N=21 cells/6 mice | 128 ± 21 | 0.40 ± 0.01 | 47 ± 1.0 | 13 ± 0.8 | 268 ± 13 | 1355 ± 101 |
| Preadolescent *Grin2a*<sup>+/-</sup><br>N=18 cells/5 mice | 135 ± 21 | 0.45 ± 0.05 | 46 ± 2 | 14 ± 0.6 | 227 ± 14 | 1083 ± 108 |
| Preadolescent *Grin2a*<sup>-/-</sup><br>N=18 cells/5 mice | 93 ± 19 | 0.54 ± 0.04 | 45 ± 3 | 11 ± 1.6 | 186 ± 18 | 761 ± 111 |
| Adult *Grin2a*<sup>+/+</sup><br>N=12 cells/6 mice | 128 ± 21 | 0.48 ± 0.02 | 39 ± 2.5 | 11 ± 1.0 | 247 ± 16 | 1150 ± 161 |
| Adult *Grin2a*<sup>+/-</sup><br>N=12 cells/4 mice | 103 ± 13 | 0.46 ± 0.03 | 46 ± 2.1 | 12 ± 0.6 | 225 ± 20 | 1145 ± 124 |
| Adult *Grin2a*<sup>-/-</sup><br>N=14 cells/5 mice | 130 ± 19 | 0.45 ± 0.02 | 38 ± 3.3 | 10 ± 0.9 | 244 ± 18 | 1161 ± 170 |

## References

Adesnik, H., G. Li, M. J. During, S. J. Pleasure, and R. A. Nicoll. 2008. ‘NMDA receptors inhibit synapse unsilencing during brain development’, Proc Natl Acad Sci U S A, 105: 5597–602.

Amin, J. B., G. R. Moody, and L. P. Wollmuth. 2021. ‘From bedside-to-bench: What disease-associated variants are teaching us about the NMDA receptor’, J Physiol, 599: 397–416.

Benke, T. A., K. Park, I. Krey, C. R. Camp, R. Song, A. J. Ramsey, H. Yuan, S. F. Traynelis, and J. Lemke. 2021. ‘Clinical and therapeutic significance of genetic variation in the GRIN gene family encoding NMDARs’, Neuropharmacology, 199: 108805.

Booker, S. A., A. Sumera, P. C. Kind, and D. J. A. Wyllie. 2021. ‘Contribution of NMDA Receptors to Synaptic Function in Rat Hippocampal Interneurons’, eNeuro, 8.

Bossi, S., D. Dhanasobhon, G. C. R. Ellis-Davies, J. Frontera, M. de Brito Van Velze, J. Lourenco, A. Murillo, R. Lujan, M. Casado, I. Perez-Otano, A. Bacci, D. Popa, P. Paoletti, and N. Rebola. 2022. ‘GluN3A excitatory glycine receptors control adult cortical and amygdalar circuits’, Neuron, 110: 2438–54 e8.

Calin, A., A. S. Ilie, and C. J. Akerman. 2021. ‘Disrupting epileptiform activity by preventing parvalbumin interneuron depolarization block’, J Neurosci.

Carmignoto, G., and S. Vicini. 1992. ‘Activity-dependent decrease in NMDA receptor responses during development of the visual cortex’, Science, 258: 1007–11.

Carvill, G. L., B. M. Regan, S. C. Yendle, B. J. O’Roak, N. Lozovaya, N. Bruneau, N. Burnashev, A. Khan, J. Cook, E. Geraghty, L. G. Sadleir, S. J. Turner, M. H. Tsai, R. Webster, R. Ouvrier, J. A. Damiano, S. F. Berkovic, J. Shendure, M. S. Hildebrand, P. Szepetowski, I. E. Scheffer, and H. C. Mefford. 2013. ‘GRIN2A mutations cause epilepsy-aphasia spectrum disorders’, Nat Genet, 45: 1073–6.

Chittajallu, R., J. C. Wester, M. T. Craig, E. Barksdale, X. Q. Yuan, G. Akgul, C. Fang, D. Collins, S. Hunt, K. A. Pelkey, and C. J. McBain. 2017. ‘Afferent specific role of NMDA receptors for the circuit integration of hippocampal neurogliaform cells’, Nat Commun, 8: 152.

Collingridge, G. 1987. ‘Synaptic plasticity. The role of NMDA receptors in learning and memory’, Nature, 330: 604–5.

de Lecea, L., J. A. del Rio, and E. Soriano. 1995. ‘Developmental expression of parvalbumin mRNA in the cerebral cortex and hippocampus of the rat’, Brain Res Mol Brain Res, 32: 1–13.

Dehorter, N., G. Ciceri, G. Bartolini, L. Lim, I. del Pino, and O. Marin. 2015. ‘Tuning of fast-spiking interneuron properties by an activity-dependent transcriptional switch’, Science, 349: 1216–20.

Denaxa, M., G. Neves, A. Rabinowitz, S. Kemlo, P. Liodis, J. Burrone, and V. Pachnis. 2018. ‘Modulation of Apoptosis Controls Inhibitory Interneuron Number in the Cortex’, Cell Rep, 22: 1710–21.

Doischer, D., J. A. Hosp, Y. Yanagawa, K. Obata, P. Jonas, I. Vida, and M. Bartos. 2008. ‘Postnatal differentiation of basket cells from slow to fast signaling devices’, J Neurosci, 28: 12956–68.

Ekins, T. G., V. Mahadevan, Y. Zhang, J. A. D’Amour, G. Akgul, T. J. Petros, and C. J. McBain. 2020. ‘Emergence of non-canonical parvalbumin-containing interneurons in hippocampus of a murine model of type I lissencephaly’, Elife, 9.

Erreger, K., S. M. Dravid, T. G. Banke, D. J. Wyllie, and S. F. Traynelis. 2005. ‘Subunit-specific gating controls rat NR1/NR2A and NR1/NR2B NMDA channel kinetics and synaptic signalling profiles’, J Physiol, 563: 345–58.

Favuzzi, E., R. Deogracias, A. Marques-Smith, P. Maeso, J. Jezequel, D. Exposito-Alonso, M. Balia, T. Kroon, A. J. Hinojosa, F. Maraver E, and B. Rico. 2019. ‘Distinct molecular programs regulate synapse specificity in cortical inhibitory circuits’, Science, 363: 413–17.

Garst-Orozco, J., R. Malik, T. A. Lanz, M. L. Weber, H. Xi, D. Arion, J. F. Enwright, 3rd, D. A. Lewis, P. O’Donnell, V. S. Sohal, and D. L. Buhl. 2020. ‘GluN2D-mediated excitatory drive onto medial prefrontal cortical PV+ fast-spiking inhibitory interneurons’, PLoS One, 15: e0233895.

Goldberg, E. M., H. Y. Jeong, I. Kruglikov, R. Tremblay, R. M. Lazarenko, and B. Rudy. 2011. ‘Rapid developmental maturation of neocortical FS cell intrinsic excitability’, Cereb Cortex, 21: 666–82.

Hansen, K. B., K. K. Ogden, H. Yuan, and S. F. Traynelis. 2014. ‘Distinct functional and pharmacological properties of Triheteromeric GluN1/GluN2A/GluN2B NMDA receptors’, Neuron, 81: 1084–96.

Hansen, K. B., L. P. Wollmuth, D. Bowie, H. Furukawa, F. S. Menniti, A. I. Sobolevsky, G. T. Swanson, S. A. Swanger, I. H. Greger, T. Nakagawa, C. J. McBain, V. Jayaraman, C. M. Low, M. L. Dell’Acqua, J. S. Diamond, C. R. Camp, R. E. Perszyk, H. Yuan, and S. F. Traynelis. 2021. ‘Structure, Function, and Pharmacology of Glutamate Receptor Ion Channels’, Pharmacol Rev, 73: 298–487.

Hanson, E., M. Armbruster, L. A. Lau, M. E. Sommer, Z. J. Klaft, S. A. Swanger, S. F. Traynelis, S. J. Moss, F. Noubary, J. Chadchankar, and C. G. Dulla. 2019. ‘Tonic Activation of GluN2C/GluN2D-Containing NMDA Receptors by Ambient Glutamate Facilitates Cortical Interneuron Maturation’, J Neurosci, 39: 3611–26.

Hanson, J. E., M. Weber, W. J. Meilandt, T. Wu, T. Luu, L. Deng, M. Shamloo, M. Sheng, K. Scearce-Levie, and Q. Zhou. 2013. ‘GluN2B antagonism affects interneurons and leads to immediate and persistent changes in synaptic plasticity, oscillations, and behavior’, Neuropsychopharmacology, 38: 1221–33.

Hensch, T. K. 2005. ‘Critical period plasticity in local cortical circuits’, Nat Rev Neurosci, 6: 877–88.

Jami, S. A., S. Cameron, J. M. Wong, E. R. Daly, A. K. McAllister, and J. A. Gray. 2021. ‘Increased excitation-inhibition balance and loss of GABAergic synapses in the serine racemase knockout model of NMDA receptor hypofunction’, J Neurophysiol, 126: 11–27.

Kaiser, T., J. T. Ting, P. Monteiro, and G. Feng. 2016. ‘Transgenic labeling of parvalbumin-expressing neurons with tdTomato’, Neuroscience, 321: 236–45.

Kannangara, T. S., B. D. Eadie, C. A. Bostrom, K. Morch, P. S. Brocardo, and B. R. Christie. 2015. ‘GluN2A-/- Mice Lack Bidirectional Synaptic Plasticity in the Dentate Gyrus and Perform Poorly on Spatial Pattern Separation Tasks’, Cereb Cortex, 25: 2102–13.

Kelsch, W., Z. Li, S. Wieland, O. Senkov, A. Herb, C. Gongrich, and H. Monyer. 2014. ‘GluN2B-containing NMDA receptors promote glutamate synapse development in hippocampal interneurons’, J Neurosci, 34: 16022–30.

Kirson, E. D., and Y. Yaari. 1996. ‘Synaptic NMDA receptors in developing mouse hippocampal neurones: functional properties and sensitivity to ifenprodil’, J Physiol, 497 (Pt 2): 437–55.

Komuro, H., and P. Rakic. 1993. ‘Modulation of neuronal migration by NMDA receptors’, Science, 260: 95–7.

Lim, L., D. Mi, A. Llorca, and O. Marin. 2018. ‘Development and Functional Diversification of Cortical Interneurons’, Neuron, 100: 294–313.

Matta, J. A., K. A. Pelkey, M. T. Craig, R. Chittajallu, B. W. Jeffries, and C. J. McBain. 2013. ‘Developmental origin dictates interneuron AMPA and NMDA receptor subunit composition and plasticity’, Nat Neurosci, 16: 1032–41.

Miller, M. N., B. W. Okaty, S. Kato, and S. B. Nelson. 2011. ‘Activity-dependent changes in the firing properties of neocortical fast-spiking interneurons in the absence of large changes in gene expression’, Dev Neurobiol, 71: 62–70.

Miyamae, T., K. Chen, D. A. Lewis, and G. Gonzalez-Burgos. 2017. ‘Distinct Physiological Maturation of Parvalbumin-Positive Neuron Subtypes in Mouse Prefrontal Cortex’, J Neurosci, 37: 4883–902.

Monyer, H., N. Burnashev, D. J. Laurie, B. Sakmann, and P. H. Seeburg. 1994. ‘Developmental and regional expression in the rat brain and functional properties of four NMDA receptors’, Neuron, 12: 529–40.

Murillo, A., A. I. Navarro, E. Puelles, Y. Zhang, T. J. Petros, and I. Perez-Otano. 2021. ‘Temporal Dynamics and Neuronal Specificity of Grin3a Expression in the Mouse Forebrain’, Cereb Cortex, 31: 1914–26.

Nakazawa, K., T. J. McHugh, M. A. Wilson, and S. Tonegawa. 2004. ‘NMDA receptors, place cells and hippocampal spatial memory’, Nat Rev Neurosci, 5: 361–72.

Nash, J. E., and J. M. Brotchie. 2000. ’A common signaling pathway for striatal NMDA and adenosine A2a receptors: implications for the treatment of Parkinson’s disease’, J Neurosci, 20: 7782–9.

Okaty, B. W., M. N. Miller, K. Sugino, C. M. Hempel, and S. B. Nelson. 2009. ‘Transcriptional and electrophysiological maturation of neocortical fast-spiking GABAergic interneurons’, J Neurosci, 29: 7040–52.

Oswald, A. M., and A. D. Reyes. 2008. ‘Maturation of intrinsic and synaptic properties of layer 2/3 pyramidal neurons in mouse auditory cortex’, J Neurophysiol, 99: 2998–3008.

Pelkey, K. A., R. Chittajallu, M. T. Craig, L. Tricoire, J. C. Wester, and C. J. McBain. 2017. ‘Hippocampal GABAergic Inhibitory Interneurons’, Physiol Rev, 97: 1619–747.

Perszyk, R. E., J. O. DiRaddo, K. L. Strong, C. M. Low, K. K. Ogden, A. Khatri, G. A. Vargish, K. A. Pelkey, L. Tricoire, D. C. Liotta, Y. Smith, C. J. McBain, and S. F. Traynelis. 2016. ‘GluN2D-Containing N-methyl-d-Aspartate Receptors Mediate Synaptic Transmission in Hippocampal Interneurons and Regulate Interneuron Activity’, Mol Pharmacol, 90: 689–702.

Perszyk, R. E., S. J. Myers, H. Yuan, A. J. Gibb, H. Furukawa, A. I. Sobolevsky, and S. F. Traynelis. 2020. ‘Hodgkin-Huxley-Katz Prize Lecture: Genetic and pharmacological control of glutamate receptor channel through a highly conserved gating motif’, J Physiol, 598: 3071–83.

Piatti, V. C., M. G. Davies-Sala, M. S. Esposito, L. A. Mongiat, M. F. Trinchero, and A. F. Schinder. 2011. ‘The timing for neuronal maturation in the adult hippocampus is modulated by local network activity’, J Neurosci, 31: 7715–28.

Priya, R., M. F. Paredes, T. Karayannis, N. Yusuf, X. Liu, X. Jaglin, I. Graef, A. Alvarez-Buylla, and G. Fishell. 2018. ‘Activity Regulates Cell Death within Cortical Interneurons through a Calcineurin-Dependent Mechanism’, Cell Rep, 22: 1695–709.

Que, L., D. Lukacsovich, W. Luo, and C. Foldy. 2021. ‘Transcriptional and morphological profiling of parvalbumin interneuron subpopulations in the mouse hippocampus’, Nat Commun, 12: 108.

Sakimura, K., T. Kutsuwada, I. Ito, T. Manabe, C. Takayama, E. Kushiya, T. Yagi, S. Aizawa, Y. Inoue, H. Sugiyama, and, et al. 1995. ‘Reduced hippocampal LTP and spatial learning in mice lacking NMDA receptor epsilon 1 subunit’, Nature, 373: 151–5.

Salmi, M., F. Del Gallo, M. Minlebaev, A. Zakharov, V. Pauly, P. Perron, A. Pons-Bennaceur, S. Corby-Pellegrino, L. Aniksztejn, P. P. Lenck-Santini, J. Epsztein, R. Khazipov, N. Burnashev, G. Bertini, and P. Szepetowski. 2019. ‘Impaired vocal communication, sleep-related discharges, and transient alteration of slow-wave sleep in developing mice lacking the GluN2A subunit of N-methyl-d-aspartate receptors’, Epilepsia, 60: 1424–37.

Sans, N., R. S. Petralia, Y. X. Wang, J. Blahos, 2nd, J. W. Hell, and R. J. Wenthold. 2000. ‘A developmental change in NMDA receptor-associated proteins at hippocampal synapses’, J Neurosci, 20: 1260-71.

Schizophrenia Working Group of the Psychiatric Genomics, Consortium. 2014. ‘Biological insights from 108 schizophrenia-associated genetic loci’, Nature, 511: 421–7.

Schneggenburger, R. 1996. ‘Simultaneous measurement of Ca2+ influx and reversal potentials in recombinant N-methyl-D-aspartate receptor channels’, Biophys J, 70: 2165–74.

Singh, T., T. Poterba, D. Curtis, H. Akil, M. Al Eissa, J. D. Barchas, N. Bass, T. B. Bigdeli, G. Breen, E. J. Bromet, P. F. Buckley, W. E. Bunney, J. Bybjerg-Grauholm, W. F. Byerley, S. B. Chapman, W. J. Chen, C. Churchhouse, N. Craddock, C. M. Cusick, L. DeLisi, S. Dodge, M. A. Escamilla, S. Eskelinen, A. H. Fanous, S. V. Faraone, A. Fiorentino, L. Francioli, S. B. Gabriel, D. Gage, S. A. Gagliano Taliun, A. Ganna, G. Genovese, D. C. Glahn, J. Grove, M. H. Hall, E. Hamalainen, H. O. Heyne, M. Holi, D. M. Hougaard, D. P. Howrigan, H. Huang, H. G. Hwu, R. S. Kahn, H. M. Kang, K. J. Karczewski, G. Kirov, J. A. Knowles, F. S. Lee, D. S. Lehrer, F. Lescai, D. Malaspina, S. R. Marder, S. A. McCarroll, A. M. McIntosh, H. Medeiros, L. Milani, C. P. Morley, D. W. Morris, P. B. Mortensen, R. M. Myers, M. Nordentoft, N. L. O’Brien, A. M. Olivares, D. Ongur, W. H. Ouwehand, D. S. Palmer, T. Paunio, D. Quested, M. H. Rapaport, E. Rees, B. Rollins, F. K. Satterstrom, A. Schatzberg, E. Scolnick, L. J. Scott, S. I. Sharp, P. Sklar, J. W. Smoller, J. L. Sobell, M. Solomonson, E. A. Stahl, C. R. Stevens, J. Suvisaari, G. Tiao, S. J. Watson, N. A. Watts, D. H. Blackwood, A. D. Borglum, B. M. Cohen, A. P. Corvin, T. Esko, N. B. Freimer, S. J. Glatt, C. M. Hultman, A. McQuillin, A. Palotie, C. N. Pato, M. T. Pato, A. E. Pulver, D. St Clair, M. T. Tsuang, M. P. Vawter, J. T. Walters, T. M. Werge, R. A. Ophoff, P. F. Sullivan, M. J. Owen, M. Boehnke, M. C. O’Donovan, B. M. Neale, and M. J. Daly. 2022. ’Rare coding variants in ten genes confer substantial risk for schizophrenia’, Nature, 604: 509-16.

Southwell, D. G., M. F. Paredes, R. P. Galvao, D. L. Jones, R. C. Froemke, J. Y. Sebe, C. Alfaro-Cervello, Y. Tang, J. M. Garcia-Verdugo, J. L. Rubenstein, S. C. Baraban, and A. Alvarez-Buylla. 2012. ‘Intrinsically determined cell death of developing cortical interneurons’, Nature, 491: 109–13.

Spear, L. P. 2000. ‘The adolescent brain and age-related behavioral manifestations’, Neurosci Biobehav Rev, 24: 417–63.

Strehlow, V., C. Rieubland, S. Gallati, S. Kim, S. J. Myers, V. Peterson, A. J. Ramsey, D. D. Teuscher, S. F. Traynelis, and J. R. Lemke. 2022. ‘Compound-heterozygous GRIN2A null variants associated with severe developmental and epileptic encephalopathy’, Epilepsia, 63: e132–e37.

Takesian, A. E., and T. K. Hensch. 2013. ‘Balancing plasticity/stability across brain development’, Prog Brain Res, 207: 3–34.

Ultanir, S. K., J. E. Kim, B. J. Hall, T. Deerinck, M. Ellisman, and A. Ghosh. 2007. ‘Regulation of spine morphology and spine density by NMDA receptor signaling in vivo’, Proc Natl Acad Sci U S A, 104: 19553–8.

Veres, J. M., G. A. Nagy, and N. Hajos. 2017. ‘Perisomatic GABAergic synapses of basket cells effectively control principal neuron activity in amygdala networks’, Elife, 6.

von Engelhardt, J., C. Bocklisch, L. Tonges, A. Herb, M. Mishina, and H. Monyer. 2015. ‘GluN2D-containing NMDA receptors-mediate synaptic currents in hippocampal interneurons and pyramidal cells in juvenile mice’, Front Cell Neurosci, 9: 95.

Wang, X. J. 2002. ‘Probabilistic decision making by slow reverberation in cortical circuits’, Neuron, 36: 955–68.

Watanabe, D., H. Inokawa, K. Hashimoto, N. Suzuki, M. Kano, R. Shigemoto, T. Hirano, K. Toyama, S. Kaneko, M. Yokoi, K. Moriyoshi, M. Suzuki, K. Kobayashi, T. Nagatsu, R. J. Kreitman, I. Pastan, and S. Nakanishi. 1998. ‘Ablation of cerebellar Golgi cells disrupts synaptic integration involving GABA inhibition and NMDA receptor activation in motor coordination’, Cell, 95: 17–27.

Williams, K., S. L. Russell, Y. M. Shen, and P. B. Molinoff. 1993. ‘Developmental switch in the expression of NMDA receptors occurs in vivo and in vitro’, Neuron, 10: 267–78.

Wong, F. K., K. Bercsenyi, V. Sreenivasan, A. Portales, M. Fernandez-Otero, and O. Marin. 2018. ‘Pyramidal cell regulation of interneuron survival sculpts cortical networks’, Nature, 557: 668–73.

Yuan, H., K. B. Hansen, K. M. Vance, K. K. Ogden, and S. F. Traynelis. 2009. ‘Control of NMDA receptor function by the NR2 subunit amino-terminal domain’, J Neurosci, 29: 12045–58.

Zhang, Z. W. 2004. ‘Maturation of layer V pyramidal neurons in the rat prefrontal cortex: intrinsic properties and synaptic function’, J Neurophysiol, 91: 1171–82.

